# A scalable, clinically severe pig model for Duchenne muscular dystrophy

**DOI:** 10.1101/2021.09.01.457562

**Authors:** Michael Stirm, Lina Marie Fonteyne, Bachuki Shashikadze, Magdalena Lindner, Maila Chirivi, Andreas Lange, Clara Kaufhold, Christian Mayer, Ivica Medugorac, Barbara Kessler, Mayuko Kurome, Valeri Zakhartchenko, Arne Hinrichs, Elisabeth Kemter, Sabine Krause, Rüdiger Wanke, Georg J. Arnold, Gerhard Wess, Hiroshi Nagashima, Martin Hrabĕ de Angelis, Florian Flenkenthaler, Levin Arne Kobelke, Claudia Bearzi, Roberto Rizzi, Andrea Bähr, Kaspar Matiasek, Maggie C. Walter, Christian Kupatt, Sibylle Ziegler, Peter Bartenstein, Thomas Fröhlich, Nikolai Klymiuk, Andreas Blutke, Eckhard Wolf

**Author notes:** Correspondence: Eckhard Wolf, Gene Center, LMU Munich, Germany. equal first author contribution (Michael Stirm, Lina Marie Fonteyne). shared last authorship (Nikolai Klymiuk, Andreas Blutke, Eckhard Wolf).

## Abstract

Large animal models for Duchenne muscular dystrophy (DMD) are crucial for preclinical evaluation of novel diagnostic procedures and treatment strategies. Pigs cloned from male cells lacking *DMD* exon 52 (*DMD*Δ52) resemble molecular, clinical and pathological hallmarks of DMD, but cannot be propagated by breeding due to death before sexual maturity. Therefore, female *DMD*^+/-^ carriers were generated. A single founder animal had 11 litters with 29 *DMD*^Y/-^, 34 *DMD*^+/-^ as well as 36 male and 29 female wild-type (WT) offspring. Breeding with F1 and F2 *DMD*^+/-^ carriers resulted in additional 114 *DMD*^Y/-^ piglets. The majority of them survived for 3-4 months, providing large cohorts for experimental studies. Pathological investigations and proteome studies of skeletal muscles and myocardium confirmed the resemblance of human disease mechanisms. Importantly, *DMD*^Y/-^ pigs reveal progressive fibrosis of myocardium and increased expression of connexin-43, associated with significantly reduced left ventricular fractional shortening and ejection fraction already at age 3 months. Furthermore, behavioral tests provided evidence for impaired cognitive ability of *DMD*^Y/-^ pigs. Our breeding cohort of *DMD*Δ52 pigs and standardized tissue repositories from *DMD*^Y/-^ pigs, *DMD*^+/-^ carriers, and WT littermate controls provide important resources for studying DMD disease mechanisms and for testing novel diagnostic procedures and treatment strategies.

## Introduction

Duchenne muscular dystrophy (DMD) is a severe, progressive, muscle-wasting disease caused by mutations in the X-linked gene *DMD* (encoding dystrophin) leading to a lack of dystrophin in muscle. Most patients lose ambulation around age 10–12 years and need assisted ventilation at around 20 years. With optimal care, most DMD patients die between 20 and 40 years of age from cardiac and/or respiratory failure (for a comprehensive introduction into DMD epidemiology, inheritance pattern and pathophysiology, see (1)).

Animal models for DMD are inevitable for studying disease mechanisms and for testing therapeutic concepts. The classical *mdx* mouse with a nonsense mutation in *Dmd* exon 23 and several other strains resembling specific human mutations are most widely used, but have limitations regarding the resemblance of the human disease phenotype. In particular, they do not develop early dilated cardiomyopathy as seen in patients (reviewed in (2)). Nevertheless, *mdx* mouse models have played key roles in studies of DMD pathogenesis and treatment development. The most widely used large animal model for DMD is the golden retriever muscular dystrophy (GRMD) dog, which has a single base change in intron 6 leading to skipping of exon 7 and the termination of the *DMD* reading frame. The animal model shows myofiber degeneration, fibrosis and fatty infiltration reflecting the histomorphological changes seen in human DMD patients. Moreover, signs of cardiomyopathy have been recorded. However, generation of large animal numbers, sufficient for relevant statistical group sizes, is limited by long generation intervals due to season breeding and relatively small litter sizes (reviewed in (3)). This also applies for other dog lines with spontaneous *DMD* mutations (summarized in (3)). Additional DMD models have been developed by gene editing in rat (4, 5), rabbit (6), and rhesus monkey (7). The specific pros and cons of these models have been reviewed recently and none of these models has so far been used for testing therapies (8). Attempts to generate a porcine DMD pig model by injection of CRISPR/Cas9 into zygotes were hampered by mosaicism and early death of the offspring (9).

Klymiuk, et al. (10) generated the first porcine DMD model. The strategy was based on targeted deletion of *DMD* exon 52 (*DMD*Δ52) by homologous recombination with a modified bacterial artificial chromosome vector in a male porcine kidney cell line and subsequent use of targeted cell clones for somatic cell nuclear transfer (SCNT). The cloned *DMD*^Y/-^ piglets lacked dystrophin and resembled clinical, biochemical and pathological hallmarks of the human disease. However birth weights of the animals were – presumably as a consequence of the SCNT procedure – highly variable and most of the animals died within the first week of life. None of the pigs reached sexual maturity; thus cloning or blastocyst complementation (11) were the only ways to propagate the model.

To overcome this limitation, we generated *DMD*^+/-^ carrier pigs, which allows scaling up the generation of *DMD*^Y/-^ pigs by breeding. In this study, we performed a comprehensive characterization of this translational animal model, revealing characteristic pathological alterations and proteome changes in skeletal muscle and myocardium, associated with impaired heart function. Moreover, we provide evidence for impaired cognitive ability of *DMD*^Y/-^ pigs. For the first time, cardiac alterations of *DMD*^+/-^ carrier pigs were characterized.

## Results

### Generation of heterozygous DMDΔ52 (DMD^+/-^) carriers and establishment of a breeding herd

A heterozygous *DMD*Δ52 mutation was introduced in female primary kidney cells (German Landrace × Swabian-Hall hybrid background) by homologous recombination with a modified bacterial artificial chromosome (BAC) in which *DMD* exon 52 was replaced by a neomycin resistance cassette (Figure 1a). The loss of exon 52 causes a shift of the reading frame, resulting in premature stop codons (Figure 1b). A correctly targeted single-cell clone was used for SCNT. Transfer of cloned *DMD*^+/-^ embryos to recipients resulted in the birth of 7 live *DMD*^+/-^ piglets, of which one (#3040; Figure 1c) could be raised to adulthood.

**Figure 1.**
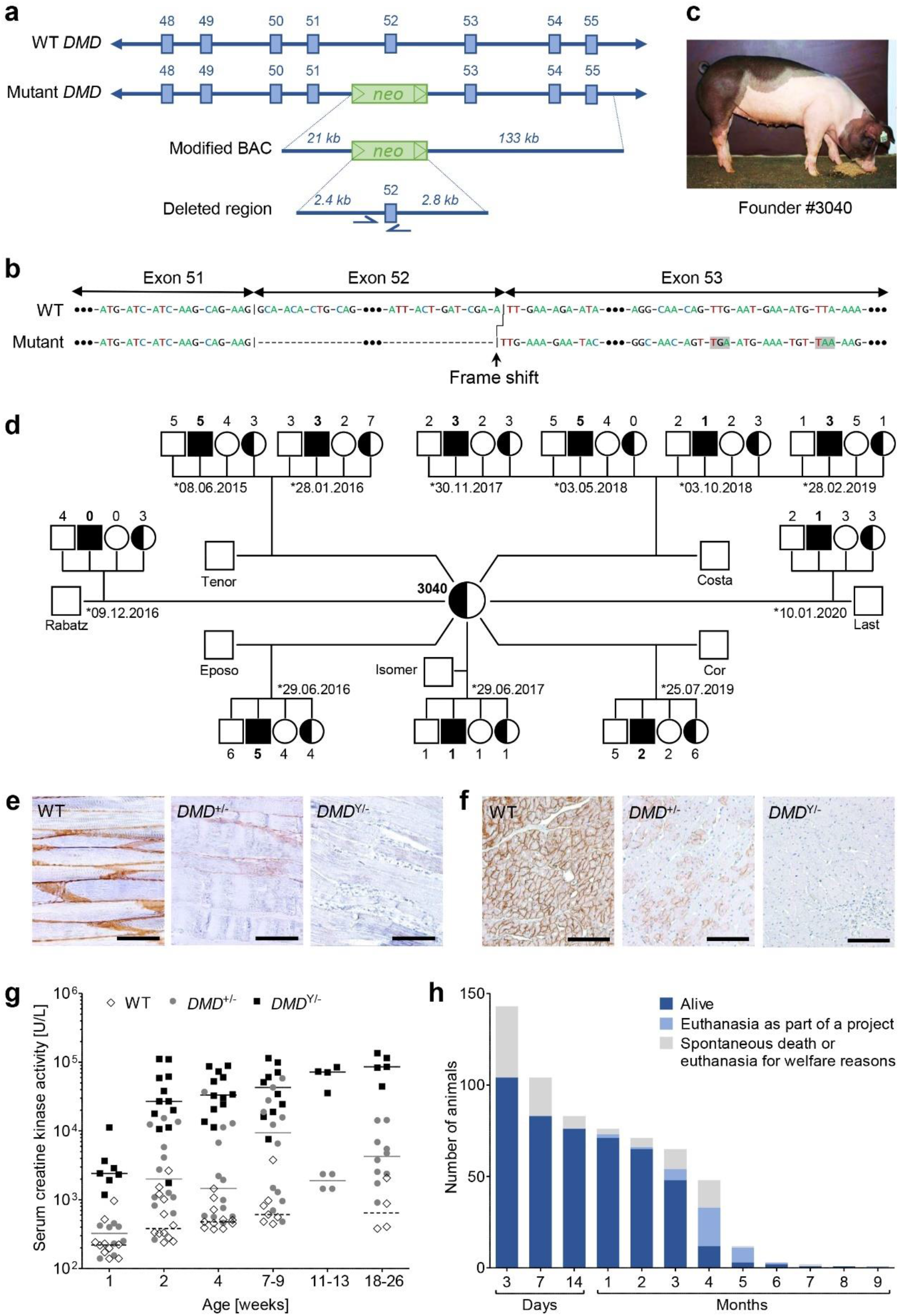
Generation of a *DMD*^+/-^ carrier and establishment of a breeding colony. **a**) Targeting strategy for the deletion of *DMD* exon 52. **b**) Frameshift and downstream premature stop codons (shaded in grey) in the cDNA caused by the loss of exon 52 of the *DMD* gene. **c**) Founder carrier sow #3040. **d**) Pedigree established by mating #3040 with wild-type boars. The total number of offspring is 29 *DMD*^Y/-^, 34 *DMD*^+/-^, 36 wild-type males (WT m), and 29 wild-type females (WT f), in line with the expected Mendelian ratios. **e**,**f**) Immunohistochemical detection of dystrophin in skeletal muscle (triceps brachii) (**e**) and myocardium (**f**) of 6-month-old animals. Paraffin sections, scale bars = 100 µm. **g**) Serum creatine kinase activities in *DMD*^Y/-^, *DMD*^+/-^ and WT pigs. **h**) Life expectancy of *DMD*^Y/-^ pigs.

To generate a breeding herd, the founder sow #3040 was inseminated with semen from 7 different wild-type (WT) boars, resulting in 11 litters with a total number of 128 piglets, between June 2015 and September 2020 (Figure 1d). Genotyping by polymerase chain reaction (PCR) tests specific for the wild-type or mutant *DMD* allele identified 29 male *DMD*-KO (*DMD*^Y/-^) and 34 female heterozygous carrier (*DMD*^+/-^) piglets as well as 36 male and 29 female WT littermates, approximating the expected Mendelian ratios. Immunostaining of skeletal muscle (Figure 1e) and myocardium (Figure 1f) sections revealed absence of dystrophin in *DMD*^Y/-^ samples. In WT pigs, a strong membranous pattern is present. In *DMD*^+/-^ carrier sows, myocyte membranes display a mosaic-like abundance pattern with generally markedly reduced staining intensity.

In addition to the founder sow #3040, 6 F1 and 3 F2 *DMD*^+/-^ sows were used for breeding and inseminated with semen from WT boars, resulting in another 114 *DMD*^Y/-^ and 118 *DMD*^+/-^ piglets as well as 98 male and 98 female WT littermates (Supplemental table 1).

The body weights of newborn *DMD*^Y/-^ piglets (1,212 ± 268 g; n = 51) were significantly (p = 0.0086) lower than those of their male WT littermates (1,373 ± 345 g; n = 54). Furthermore, the *DMD*^Y/-^ piglets lost weight during the first 24 hours, whereas their WT siblings gained weight. Subsequently, also the *DMD*^Y/-^ piglets gained weight but never reached the body weight of their male WT littermates (Supplemental figure 1).

Serum creatine kinase (CK) activties were highly elevated in *DMD*^Y/-^ compared to WT pigs. In *DMD*^+/-^ pigs, a wide variation in serum CK activities was observed, ranging from levels within the cluster of WT pigs to highly elevated levels as in *DMD*^Y/-^ pigs, similar to human dystrophinopathy carriers (12) (Figure 1g).

The life expectancy of *DMD*^Y/-^ pigs was significantly reduced. Of the total of n = 143 *DMD*^Y/-^ animals, 42% (n = 60) died within the first week and another 8% (n = 12) within the first month, whereas 48% (n = 68) survived between > 1 month and 5 months. So far, 3 *DMD*^Y/-^ boars survived for more than 5 months (#7259: 177 days; #6964: 192 days; #6790: 266 days) (Figure 1h). Through improved management, neonatal lethality could be reduced to 23% (12/52; Supplemental figure 2).

### Mapping of chromosome regions influencing the life expectancy of DMD^Y/-^ pigs

Since genetic modifiers have been shown to influence the clinical severity of DMD (reviewed in (13, 14)), we used our large cohort of *DMD*^Y/-^ pigs to map chromosome regions influencing life expectancy. A combined linkage disequilibrium and linkage analysis (cLDLA) with single-nucleotide polymorphism (SNP) genotypes (60k) was performed using DNA samples from 98 *DMD*^Y/-^ pigs, which had to be euthanized in a terminal condition or died spontaneously. Life expectancy was considered as continuous variable (in days). The quantitative model revealed significant chromosome-wide associations on chromosomes 6 (around 32.2 Mbp), 8 (around 33.2 Mbp), and 15 (around 100.4 Mbp) (Supplemental figure 3a). The association between haplotypes and life expectancy in the corresponding chromosome regions and associated genes are shown in Supplemental figure 3b-d.

### Age-related pathological alterations of skeletal muscles and myocardium of DMD^Y/-^ pigs

All *DMD*^Y/-^ piglets that died within the first week displayed severe gross and histopathological skeletal muscle lesion patterns, consistent with the alterations found in cloned dystrophin-deficient piglets (10). The diffusely pale skeletal musculature showed numerous foci of white, streaky, muscle necroses over the entire length of the muscle bundle (Supplemental figure 4a). Muscle fiber cross section profiles displayed a reduced size (as compared to WT sections) with marked size variation, presence of numerous rounded fibers and fibers with centralized nuclei, as well as multiple, often coalescing foci of degenerating and necrotic fibers with loss of cross-striation, hyalinization and fragmentation of the sarcoplasm, rupture and loss of fibers, and replacement by homogenous cellular and karyorrhectic debris (Figure 2a). These alterations were accompanied by interstitial edema, infiltration of inflammatory cells, including macrophages, eosinophil granulocytes, few lymphocytes and plasma cells as well as muscle fiber regeneration with hypertrophic basophilic fibers and hyperplastic satellite cells. Gross- or histomorphological myocardial lesions were not observed in this age-group (Figure 2b).

**Figure 2.**
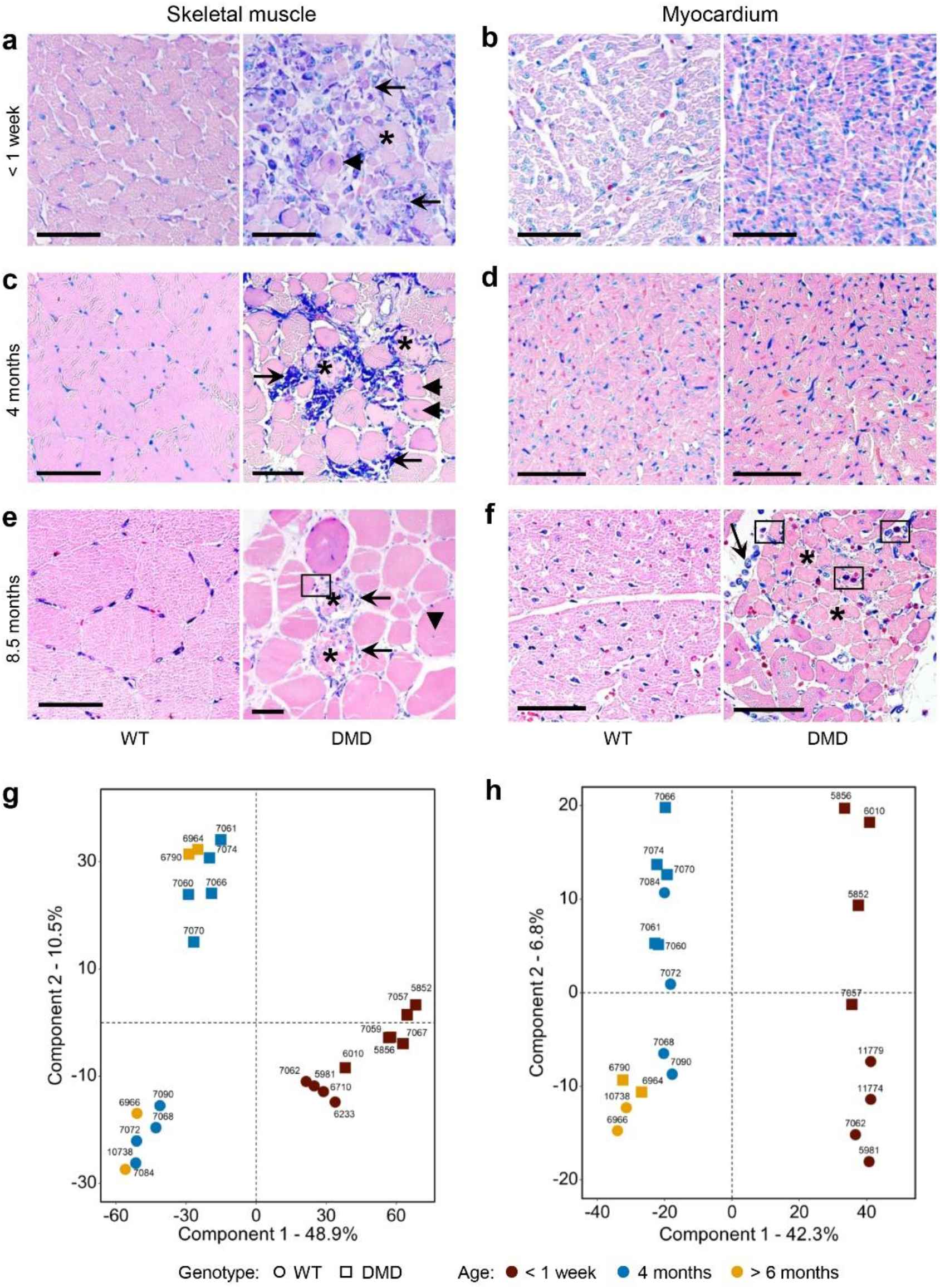
Histological changes and clustering of proteome profiles of skeletal muscle and heart from *DMD*^Y/-^ pigs surviving less than one week (**a**,**b**), 3-4 months (**c**,**d**) or more than 6 months (**e**,**f**). **a**) Histology of skeletal muscles from short-term survivors showed inflammatory cell infiltration (arrows) comprising of macrophages/histiocytes, plasma cells and few lymphocytes, degenerating muscle cells with centralized nuclei (arrowhead), and necrotic myocytes (asterisk). Myocardial sections did not display histopathological alterations evident on the light-microscopic level. **c**) In *DMD*^Y/-^ pigs surviving for 3-4 months, multiple skeletal muscles showed a similar spectrum of alterations as in a), but the lesions were more pronounced and advanced. **d**) Myocardium did not show prominent changes at the light-microscopic level. **e**) Skeletal muscle from the longest surviving *DMD*^Y/-^ boar #6790 displayed the changes described above accompanied by infiltration of eosinophilic granulocytes (rectangle). At this age, obvious histological alterations of myocardium were revealed. Histology: Paraffin sections, Giemsa-staining. Bars = 50 µm. **g**,**h**) Principal component analyses of proteome profiles from skeletal muscle (**g**) and myocardium (**h**) tissue samples of *DMD*^Y/-^and WT pigs of different age groups.

Corresponding macroscopic (Supplemental figure 4b) and microscopic (Figure 2c) skeletal muscle lesion patterns were also regularly identified in *DMD*^Y/-^ pigs surviving between 1 and 5 months; however, foci of necrotic muscle fibers and corresponding inflammatory alterations were markedly less frequent and severe compared to the piglets that died within the first week. Significant myocardial lesions were also not observed in this age-cohort (Figure 2d).

From 4-month-old *DMD*^Y/-^ pigs (n = 6) and WT littermates (n = 4) a complex biobank including almost 40 different tissues and body fluids was established for morphological and molecular profiling studies (Supplemental table 2).

Two long-term surviving *DMD*^Y/-^ boars deserve particular attention. #6790 (Supplemental figure 4c) was trained to mount a dummy sow from 6 months of age. However, the collection of semen was not successful. After showing first signs of stenotic breathing/limited respiratory function (breathing through the mouth, snoring, shortness of breath worsening on excitation or lying down, cough), #6790 was euthanized for sampling at an age of 9 months (266 days). At this time, his body weight (76,5 kg) was severly reduced compared to WT (142 kg at 6.5 months; see below). Necropsy revealed myocardial alterations, grossly presenting as pale, white-to-tan, indistinctly limited, subepicardial and intramyocardial foci of few mm diameter (Supplemental figure 4c, macroscopic imaging of the heart). #6964 was successfully trained to mount a dummy sow for the collection of semen, which was used for the insemination of 4 *DMD*^+/-^ sows. The founder sow #3040 and an F1 sow (#5382) became pregnant and gave birth to a total of n = 28 live piglets (15 male WT, 4 *DMD*^+/-^, 5 *DMD*^-/-^, 5 *DMD*^Y/-^) (Supplemental figure 4d). Boar #6964 died peracutely at age 192 days, immediately after successful semen collection, most probably due to sudden cardiac death. The animal showed a greatly reduced body weight (93.5 kg) with normal body condition, when compared to a male WT sibling (142 kg) raised under the same conditions.

Both long-term surviving *DMD*^Y/-^ boars exhibited the characteristic histopathological skeletal muscle fiber alteration patterns associated with DMD deficiency, with occasional foci of necrosis and histiocytic inflammation with participation of eosinophils (Figure 2e). The myocardial foci of #6790 consisted of degenerating and necrotic cardiomyocytes with interstitial edema, and infiltration of macrophages, accompanied by considerable numbers of eosinophils (Figure 2f). Although #6964 did not display macroscopic alterations of the heart, histopathologic alterations, reflecting the situation as in #6790, were observed (data not shown).

### Age-related alterations of skeletal muscle and myocardial proteome profiles in DMD^Y/-^ pigs

To analyze molecular alterations associated with the pathological changes of skeletal muscle and myocardium in the different age groups of *DMD*^Y/-^ pigs label-free proteome analyses were performed. A total of 3,030 and 3,136 proteins were identified with high confidence (FDR < 0.01) in skeletal muscle and myocardium, respectively. A comprehensive description of the results is provided in Supplemental table 3. The data set has been submitted to the ProteomeXchange Consortium via the PRIDE (15) partner repository, http://proteomecentral.proteomexchange.org; PXD027772.

Principal component analyses (PCA) (Figure 2g,h) as well as hierachical clustering (Supplemental figure 5) of normalized protein intensity values clearly seperate proteome profiles of young (< 1 week age group) and older animals (4 months and > 6 months) of skeletal muscle and myocardium. For skeletal muscle cluster analysis shows a clear separation according to genotype (*DMD*^Y/-^, WT), which is more pronounced in the older animals. For myocardium separation is not as pronounced. This finding reflects the fact that in DMD skeletal muscle is the most affected type of muscle.

Further statistical analysis revealed 709 differentially abundant proteins in the skeletal muscle of the < 1 week age group (n = 4 for WT and n = 6 for *DMD*^Y/-^), 475 with increased and 234 with reduced abundance in *DMD*^Y/-^ vs. WT samples (Figure 3a). At age 4 months (n = 4 for WT and n = 5 for *DMD*^Y/-^), 285 differentially abundant proteins were detected, 264 with increased and 21 with reduced abundance in *DMD*^Y/-^ vs. WT samples (Figure 3b).

**Figure 3.**
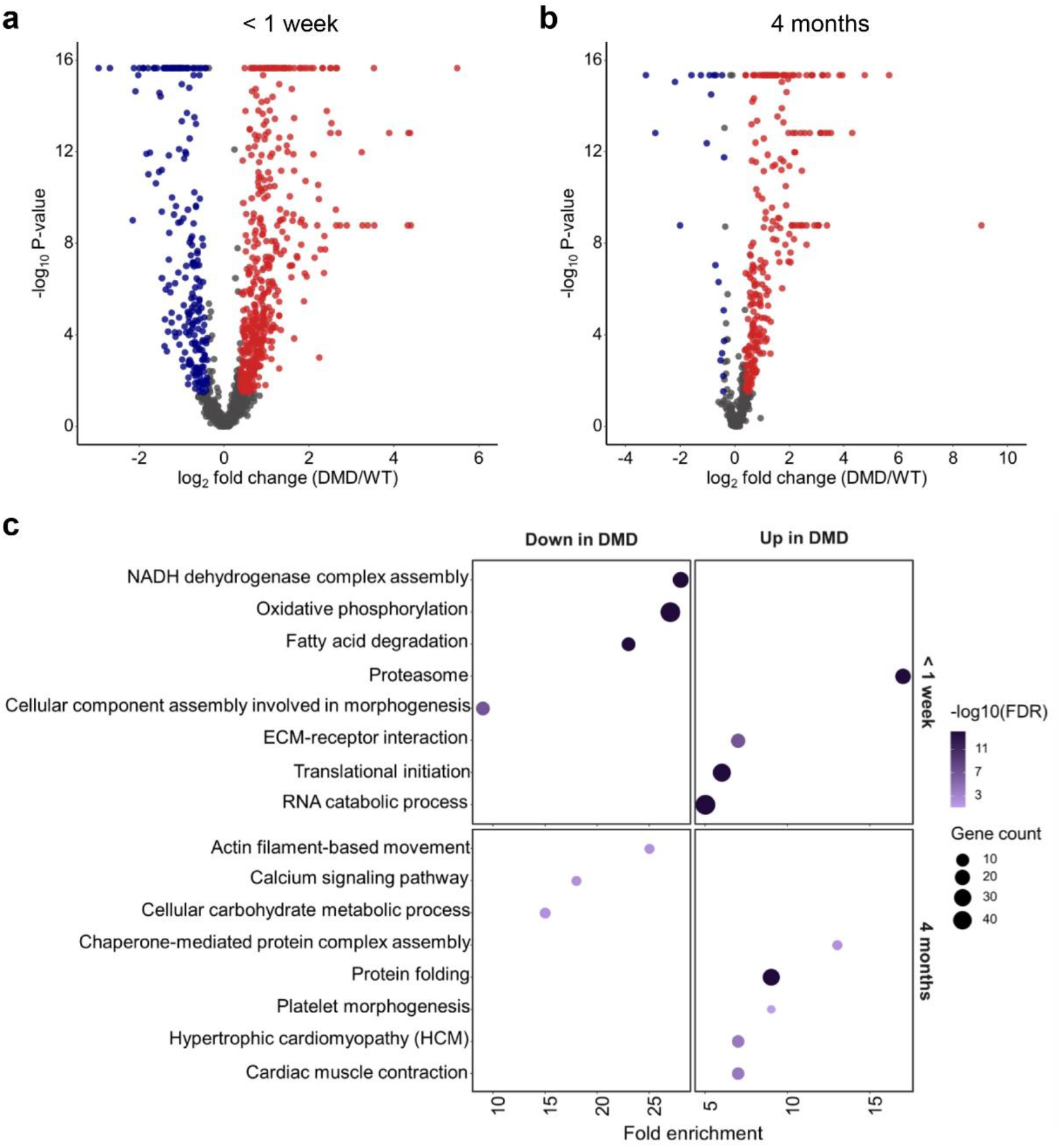
Proteome analysis of skeletal muscle. **a**,**b**) Volcano plot visualization of proteome alterations in skeletal muscle from the < 1 week age group (**a**) and the 4 month age group (**b**). Proteins significantly altered in abundance (Benjamini-Hochberg (BH) corrected p-value ≤ 0.05, ≥ 1.30- or ≤ 0.77-fold change) in *DMD*^Y/-^ are colored in blue and red for downregulation and upregulation respectively. **c**) Overrepresentation analysis using WebGestalt with gene sets according to KEGG and GO biological process databases, of skeletal muscle proteins less abundant in *DMD*^Y/-^ (left) and more abundant in *DMD*^Y/-^ (right) for < 1 week (top) and 4 month (bottom) age group. BH method was used for multiple testing adjustment. Size of the bubble indicates the corresponding number of differentially abundant proteins (referred to as gene count in the figure) and color the significance of enrichment. Fold enrichment represents magnitude of overrepresentation.

Functional annonation clustering (Figure 3c) of the differentially abundant proteins showed that in the *DMD*^Y/-^ piglets that died within the first week, proteins related to proteasome, ECM-receptor interaction, translational initiation, and RNA catabolic process were more abundant, whereas proteins involved in NADH dehydrogenase complex assembly, oxidative phosphorylation, fatty acid degradation, and morphogenesis were less abundant than in age-matched WT piglets.

The samples from 4-month-old *DMD*^Y/-^ pigs revealed increased levels of proteins related to protein folding, platelet morphogenesis, hypertrophic cardiomyopathy, and cardiac muscle contraction, whereas proteins involved in actin filament-based movement, calcium signaling, and cellular carbohydrate metabolism were decreased.

In myocardium samples from the < 1 week age group (n = 4 for WT and n = 4 for *DMD*^Y/-^), 298 differentially abundant proteins were detected, 87 with increased and 211 with reduced abundance in *DMD*^Y/-^ vs. WT samples (Figure 4a). In myocardium samples from 4-month-old animals (n = 4 for WT and n = 5 for *DMD*^Y/-^), 51 differentially abundant proteins were identified, 12 with increased and 39 with reduced abundance in *DMD*^Y/-^ vs. WT samples (Figure 4b).

**Figure 4.**
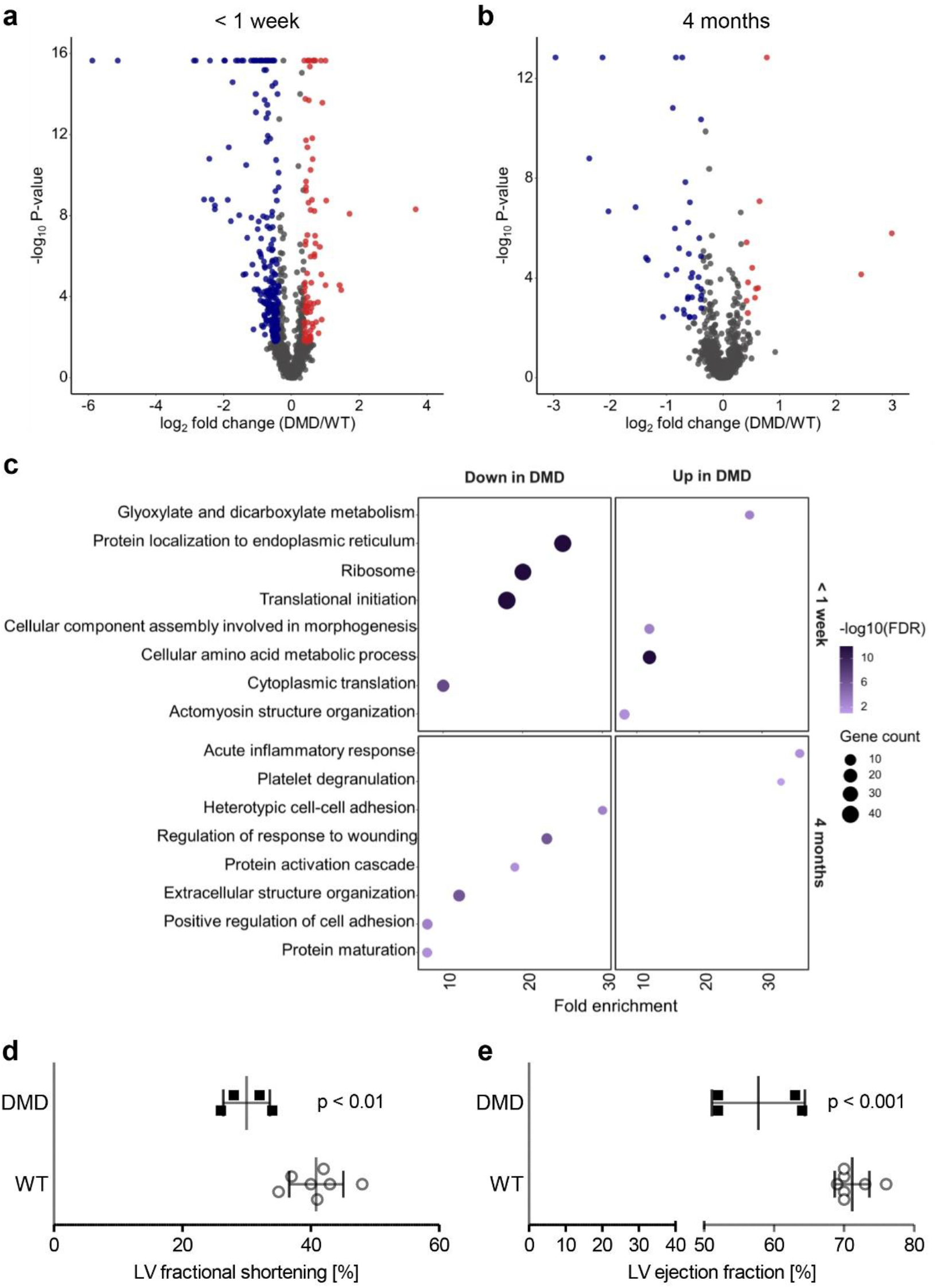
Proteome analysis of myocardium and parameters of cardiac function. **a**,**b**) Volcano plot visualization of proteome alterations in myocardium from the < 1 week age group (**a**) and the 4 month age group (**b**). Proteins significantly altered in abundance (Benjamini-Hochberg (BH) corrected p-value ≤ 0.05, ≥ 1.30- or ≤ 0.77-fold change) in *DMD*^Y/-^ are colored in blue and red for downregulation and upregulation, respectively. **c**) Overrepresentation analysis using WebGestalt with gene sets according to KEGG and GO biological process databases, of myocardium proteins less abundant in *DMD*^Y/-^ (left) and more abundant in *DMD*^Y/-^ (right) for < 1 week (top) and 4 month (bottom) age group. BH method was used for multiple testing adjustment. Size of the bubble indicates the corresponding number of differentially abundant proteins (referred to as gene count in the figure) and color the significance of enrichment. Fold enrichment represents magnitude of overrepresentation. **d**) Fractional shortening of left ventricular (LV) myocardium. **e**) LV ejection fraction.

Functional clusters enriched in *DMD*^Y/-^ piglets from the < 1 week age group were glyoxylate and dicarboxylate metabolism, cellular amino acid metabolic process, actomyosin structure organization, and cellular component assembly involved in morphogenesis. Proteins belonging to the clusters ribosome, translation, and protein localization to endoplasmic reticulum were less abundant in *DMD*^Y/-^ vs. WT samples (Figure 4c).

Functional clustering of proteins with increased abundance in 4-month-old *DMD*^Y/-^ pigs revealed the terms acute inflammatory response and platelet degranulation. Proteins with decreased abundance in *DMD*^Y/-^ vs. WT samples belong to the functional clusters cell adhesion, regulation of response to wounding, protein maturation, protein activation, and extracellular structure organization (Figure 4c).

### Impaired cardiac function in 4-month-old DMD^Y/-^ pigs

To evaluate if the proteome changes observed in myocardium are associated with impairments in cardiac function, we performed echocardiographic investigations of 4-month-old *DMD*^Y/-^ pigs (n = 4) and age-matched WT controls (n = 7). These studies revealed a 27% reduced left ventricular (LV) fractional shortening (p < 0.01; Figure 4d) and a 19% reduced LV ejection fraction (p < 0.001; Figure 4e) in *DMD*^Y/-^ compared to WT pigs. No significant dilation of the heart, decrease in the septal or parietal wall thickness were observed at this stage.

### Fibrosis and increased connexin-43 levels in DMD^Y/-^ myocardium

Since fibrosis is an early feature of cardiomyopathy in DMD (16), we evaluated the level of fibrosis in Masson trichrome-stained myocardium sections from *DMD*^Y/-^ and WT pigs of different age groups (Figure 5a). While at age < 1 week the fibrotic area was not different between the groups, it was significantly increased in *DMD*^Y/-^ pigs at age 4 months and increased further in the 6-month age group (Figure 5b).

**Figure 5.**
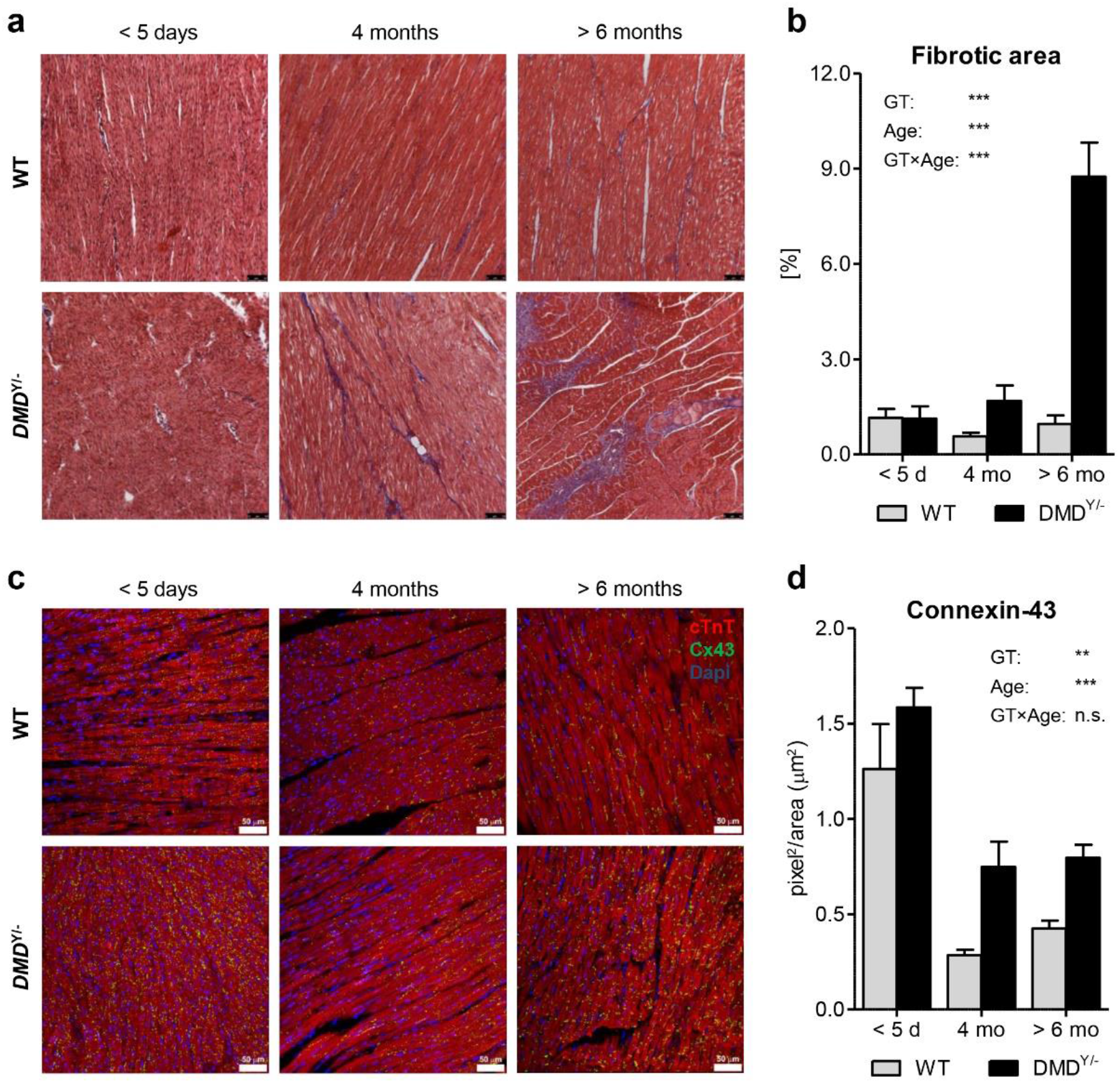
Myocardial fibrosis and connexin-43 (CX43) expression. **a**) Masson trichrome stained myocardium sections of *DMD*^Y/-^ (n = 3, 3, 2) and WT pigs (n = 3, 3, 3) at age < 5 days, 4 months, and > 6 months. Bars = 50 µm. **b**) Percentage of fibrotic area. **c**) Immunofluorescence staining for CX43 (green) and cardiac troponin T (cTnT; red) and DAPI staining (blue) of myocardial sections of *DMD*^Y/-^ and WT pigs of the three age groups. Bars = 50 µm. **d**) Quantification of CX43 in immunostained myocardium. Effects of genotype (GT), Age, and the interaction GT×Age were tested by analysis of variance. **p<0.01; ***p <0.001; n.s. = not significant.

Another aspect is the expression of connexin-43 (CX43 alias gap junction alpha-1 protein, GJA1), which has recently been identified as a factor involved in dystrophic cardiomyopathy (17). Immunofluorescence staining of myocardium sections revealed higher level of CX43 in the youngest age group than in 4- or 6-month-old animals. In the latter two age groups, the levels of CX43 were higher in *DMD*^Y/-^ than in WT pigs (Figure 5c,d).

### Gastrointestinal changes in a proportion of DMD^Y/-^ pigs

Since gastrointestinal dysfunction including constipation has been observed in patients with later stage DMD (18), this aspect was specifically investigated in 28 *DMD*^Y/-^ piglets from 9 litters. Eleven *DMD*^Y/-^ piglets were registered with difficulties in defecation (repeated strain to eliminate feces), increasing abdominal girth, anorexia, or abdominal edema and ascites (detected by ultrasound scan) within the first weeks of life. Necropsy of animals euthanized after showing either one or more of the described conditions revealed accumulation of gas and ingesta in the jejunum and large intestine, associated with obstipation, dark coloration of the intestinal wall, infarction of the intestinal wall and/or ascites and fibrinous layers in the abdominal cavity. In case of complete ileus, present in 2 animals, the intestinal wall showed porosity, with the subsequent part of the intestine being completely empty (Supplemental figure 6a). In addition to the investigation of neonatal piglets, histomorphological evaluation of the gastrointestinal tract (GIT; esophagus, gastric cardia, fundus, and pylorus, duodenum, jejunum, ileum, caecum, ascending colon, descending colon, rectum) was performed in 4-month-old *DMD*^Y/-^ piglets (n = 6) and WT siblings (n = 4). After staining with Masson’s trichrome, sections of the different parts of the GIT (gastric cardia as an example in Supplemental figure 6b) were evaluated for the presence of type I collagen fibers and scored with a system ranging from 1 to 5 (absent, sub-significant, mild, moderate, severe). Significantly (p < 0.05) increased collagen in the *DMD*^Y/-^ pigs was revealed in sections of gastric cardia and pylorus as well as caecum (Supplemental figure 6c).

### Cognitive impairment of DMD^Y/-^ pigs

To evaluate potential cognitive impairment of *DMD*^Y/-^ pigs (n = 12), we performed two behavioral tests in comparison with male littermate WT pigs (n = 13).

In the Novel Object Recognition Test, the exploration times of a known (T_old_) and a novel object (T_new_) were recorded over a period of 5 minutes. In all three replicates of the test, the total exploration time (T_total_ = T_new_ + T_old_) was significantly (p = 0.0237) shorter for the *DMD*^Y/-^ than the WT group (Figure 6a). The proportion of time spent on the new object (on average 68% and 65% for *DMD*^Y/-^ and WT) was not different between the two groups (data not shown).

**Figure 6.**
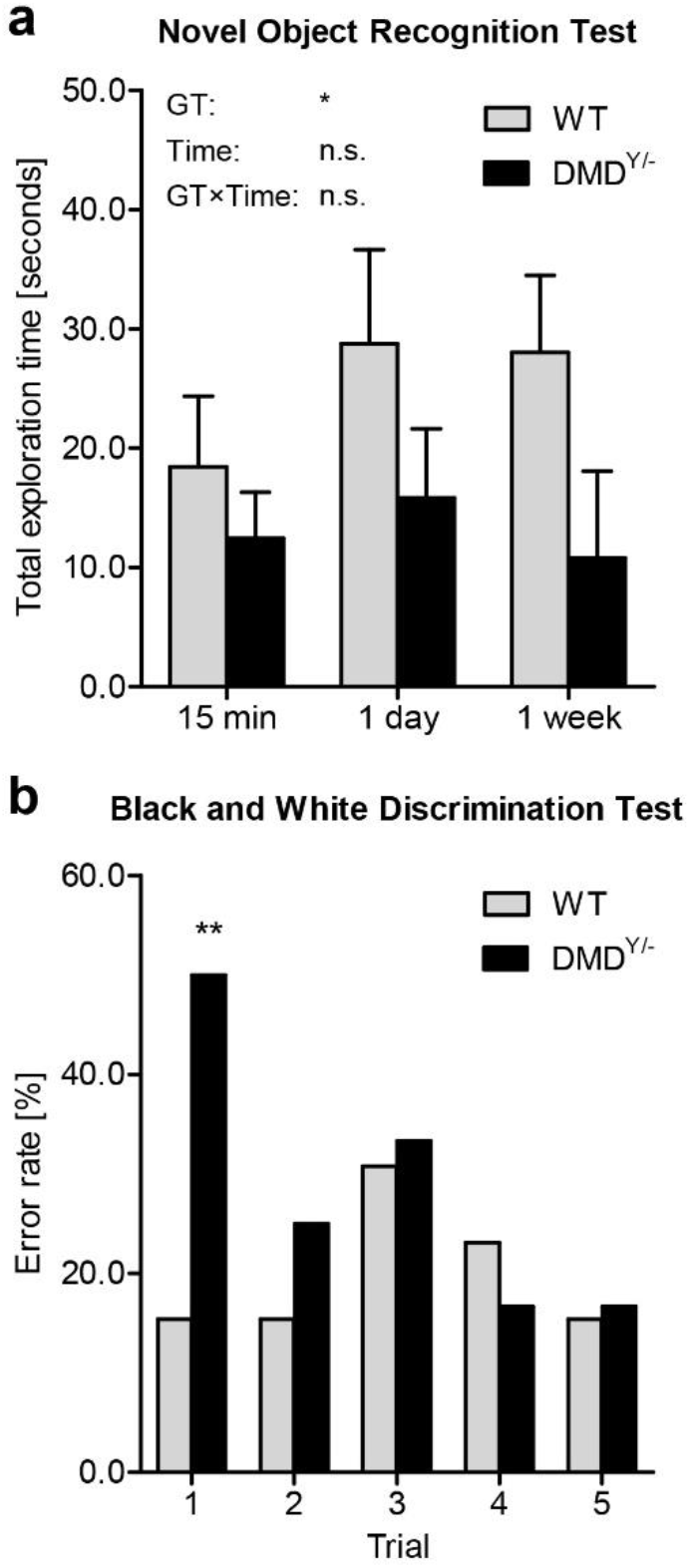
Cognitive impairment in *DMD*^Y/-^ pigs. **a**) Novel Object Recognition Test: Total time (in sec) pigs explored one of two objects (one known, one new) within a 5-min observation period in the trial box. The graph shows means and standard deviations for *DMD*^Y/-^ (n = 12) and WT pigs (n = 13) in three replicates of the test. Analysis of variance showed a significant effect of genotype (GT; *p = 0.0237), while replicate (Time) and the interaction GT×Time had no significant effects (n.s.). The proportion of time spent on the new object was not different between the two groups. **b**) Black and White Discrimination Test: Increased error rate in *DMD*^Y/-^ (n = 12) vs. WT pigs (n = 13) in the first trial after two training sessions (chi-square test: **p = 0.0089).

In a Black and White Discrimination Test, the animals had access to a reward only via one of two doors labeled with a black or white spot, respectively. In the first trial round (following two training rounds), 6/12 *DMD*^Y/-^ pigs but only 2/13 WT pigs failed (p = 0.0089). In the subsequent 4 trials, there was no difference between the two groups (Figure 6b). In a sixth round, where the positions of both doors were switched, all *DMD*^Y/-^ and WT pigs chose the wrong door (data not shown).

These observations provide evidence for at least moderate cognitive impairment of *DMD*^Y/-^ pigs, mirroring the human phenotype (19).

### Phenotypic alterations in DMD^+/-^ carrier pigs

The incidence of skeletal muscle damage among female carriers of loss-of-function *DMD* mutations, including asymptomatic carriers, was reported between 2.5% and 19%, and the incidence of dilated cardiomyopathy between 7.3% and 16.7% for DMD patients (reviewed in (12)). Therefore we took advantage of the large number of *DMD*^+/-^ animals in our pedigree to evaluate pathological lesions in skeletal muscles and myocardium.

The *DMD*^+/-^ founder sow #3040, who was clinically healthy throughout her life and did not show clinical evidence for muscle damage, was euthanized at an age of almost 7 years. Macroscopically, the heart showed severe dilation of the right ventricle associated with white, patchy discoloration of the epicard extending into the myocardium. The endocard displayed mild fibrosis (Figure 7a). Histology showed severe myocardial infiltration of adipocytes (replacement lipomatosis), fibrosis, and multifocal myocyte degeneration with inflammatory cell infiltrates consisting of eosinophils, macrophages/histiocytes, few lymphocytes and fibroblasts (Figure 7b).

**Figure 7.**
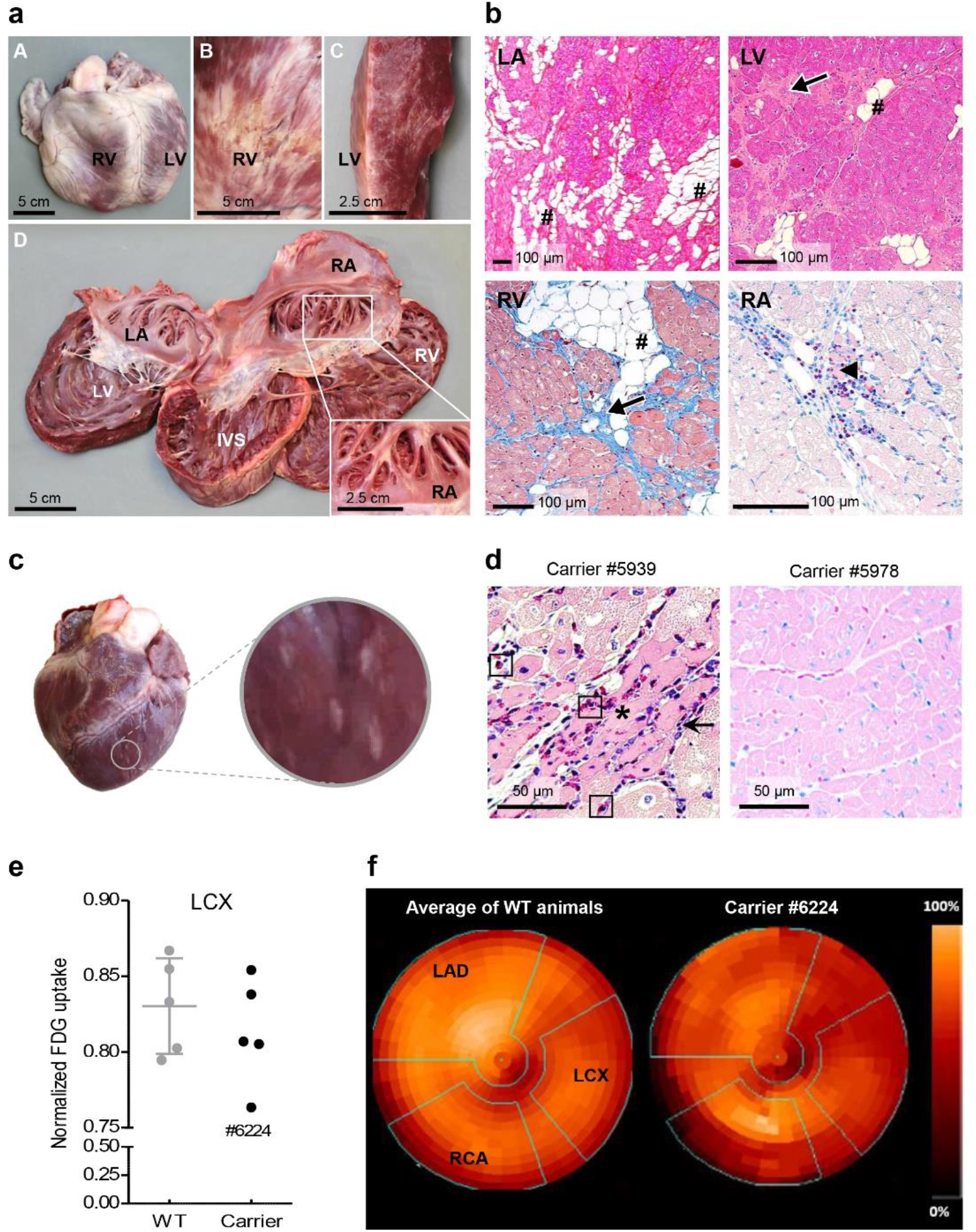
Cardiac alterations in *DMD*^+/-^ carrier pigs. **a**) Heart of 7-year-old founder sow #3040. Locations are indicated (RA: right atrium; LA: left atrium; LV: left ventricle; RV: right ventricle; IVS: interventricular septum; Ao: aorta. Note the severe dilation of the RV in A, the white, patchy discoloration of the epicard (B) that extends into the myocardium (C) and corresponds to interstitial replacement lipomatosis and fibrosis, as confirmed by histology. The endocard displays mild fibrosis (D). Note the thinning of the right and left ventricle myocardial wall. The inset to D shows a white discoloration of the endocard of the right atrial appendix. Scale bar sizes are indicated. **b**) Heart histology (different locations and stainings as indicated) displaying severe infiltration of adipocytes (replacement lipomatosis, #), fibrosis (arrows), and multifocal myocyte degeneration (arrowhead) with inflammatory cell infiltrates consisting of eosinophils, macrophages/histiocytes, few lymphocytes and fibroblasts. FFPE sections, stainings: H&E (LA, LV), Masson trichrome (RV), Giemsa (RA). Scale bar sizes are indicated. **c**) Macroscopic alteration of the heart in a 6-month-old *DMD*^+/-^ carrier pig. **d**) Variable histological alterations of skeletal muscle (triceps brachii) and mycardium in 6-month-old *DMD*^+/-^ carrier pigs. Histopathological alterations are indicated, if present (arrows: inflammatory cell infiltration comprising of macrophages/histiocytes, plasma cells and few lymphocytes; arrowheads: degenerating skeletal myocytes with centralized nuclei; asterisks: necrotic myocytes; rectangles: eosinohilic granulocytes). Paraffin sections, Giemsa-staining. Bars = 50 µm. **e**) Determination of myocardial glucose utilization by positron emission tomography/computed tomography (PET/CT) using 2-deoxy-2-[^18^F]fluoro-D-glucose (FDG) as tracer. Normalized uptake of FDG in the territory of the left circumflex artery (LCX). Note the reduced uptake in one *DMD*^+/-^ carrier animal (#6224). **f**) Polar plot representation of FDG uptake in the myocardium. Coronary artery territories are indicated by green lines (RCA = right coronary artery, LAD = left anterior descending).

Since in most symptomatic human carriers alterations become apparent around puberty (reviewed in (20)), we performed systematic necropsies of 6-month-old *DMD*^+/-^ pigs (n = 20) and female WT siblings (n = 5). The majority of the examined carrier pigs did not show macroscopic alterations of skeletal muscles or heart, but 3 animals (15%) showed prominent alterations, such as multiple white foci of myocardial necrosis and inflammation (Figure 7c). Histomorphological alterations of heart (Figure 7d) and skeletal muscles (Supplemental figure 7) were scattered within the normal tissue structure, corresponding to the areas lacking dystrophin expression. These histological changes were exclusively observed in *DMD*^+/-^ pigs, but in none of the examined WT animals. From this collective, a biobank containing samples from different skeletel muscles, different regions of the heart as well as serum and plasma samples was established (Supplemental table 4) and is available as a resource, e.g. for biomarker screening.

Next we tested if myocardial alterations can be detected at an earlier stage using a noninvasive procedure. Therefore, regional myocardial glucose uptake was determined by positron emission tomography/computed tomography (PET/CT) with the glucose-analog 2-deoxy-2-[^18^F]fluoro-D-glucose (FDG) in 3-month-old *DMD*^+/-^ pigs (n = 5) and age-matched WT controls (n = 5). Whereas the myocardial PET signals of most carriers ranged within the WT group, carrier #6224 revealed a distinctly reduced signal in the area of the left circumflex coronary artery (Figure 7e,f).

## Discussion

In this study, we developed a scalable pig model for Duchenne muscular dystrophy. The model represents one of the most common human *DMD* mutations (deletion of exon 52) and strikingly mirrors the clinical, biochemical and pathological features of the human disease in an accelerated mode. Therefore the model is highly suitable for testing new treatment strategies. Our initial model was based on *DMD* exon 52 deficient male cells, which were used to generate *DMD*^Y/-^ pigs by SCNT (10). Although the model showed the clinical phenotype and provided new insights into the molecular DMD pathology in skeletal muscle (21) and heart (22), the affected animals died before sexual maturity and could only be propagated by cloning. Epigenetic abnormalities associated with this biotechnology may per se affect the phenotype of the offspring, which was evident by a highly variable birth weight (10). One possibility to overcome this limitation is blastocyst complementation, i.e. the generation of male-female chimeras where the DMD phenotype is rescued by *DMD* intact female cells, whereas the germline is male and allows transmisson of the *DMD* mutation to the next generation (11).

Here, we generated a female heterozygous *DMD* mutant founder sow, which allowed scaling up of the model by breeding. The founder sow #3040 was clinically healthy throughout life and allowed breeding of a remarkable number of male *DMD*^Y/-^ offspring, but also of female *DMD*^+/-^ carriers and WT offspring of the same genetic background. Propagation of the porcine DMD model by breeding provides the most appropriate WT controls and also allows studies of clinical and molecular features of *DMD*^+/-^ carrier animals.

The clinical phenotype of *DMD*^Y/-^ pigs produced by breeding largely represented the phenotype of *DMD*^Y/-^ pigs produced by cloning, however any interference of the cloning technology could be excluded. In particular, piglets with a very high or low body weight were not observed. A smaller proportion of the *DMD*^Y/-^ offspring showed an early postnatal, highly severe phenotype, limiting life expectancy to less than one week. With an improved management of the piglets, the proportion of early dying animals could be reduced. The majority of *DMD*^Y/-^ pigs had a life expectancy of 3-4 months, which is very suitable for testing new therapies, since life expectancy is a stringent readout of efficacy. Only rarely (n = 3 out of 143), *DMD*^Y/-^ pigs reached a life expectancy of 6 months and beyond.

In order to identify genetic determinants of life expectancy in *DMD*^Y/-^ pigs, we performed a genome-wide association study and identified 3 regions on chromosomes 6, 8, and 15 representing quantitative trait loci with chromosome-wide significance. These regions do not contain genes whose variants are known to affect disease progression in DMD, such as osteopontin (secreted phosphoprotein 1, *SPP1*), latent TGFB protein 4 (*LTBP4*) (reviewed in (23)), and Tctex1 domain containing 1 (*TCTEX1D1* alias *DYNLT5*) (24) in humans or *JAGGED1* in dogs (25). However, our mapping study identified specific haplotypes at the QTL region of chromosome 6 with positive or negative effects on life expectancy (Supplemental table 5). These can be used for future selective breeding in order to further consolidate the phenotype of the porcine DMD model.

The severe macroscopic and histological alterations of skeletal muscles in *DMD*^Y/-^ piglets that died within the first week were associated with marked changes of the proteome, with more than 700 differentially abundant proteins compared to age-matched WT samples. The major functional classes of proteins with increased or decreased abundance are in line with muscle degeneration, regeneration and impaired metabolic activity. Although no macroscopic or microscopic alterations of the heart were observed, the proteome of the myocardium showed substantial derangements with almost 300 differentially abundant proteins.

Similarly, a larger number of proteome changes were detected in skeletal muscle of 4-month-old *DMD*^Y/-^ pigs, while proteomic derangements were less pronounced in myocardium. Overall, in skeletal muscle and myocardium differentially expressed proteins and associated biological processes and pathways largely overlap with previous observations made in cloned *DMD*^Y/-^ pigs (21, 22).

An important aspect of the porcine DMD model is cardiac involvement as shown by altered myocardial proteome profiles and in particular progressively increased fibrosis and elevated levels of connexin-43 (CX43), the major ventricular gap junction protein. Increased CX43 protein content and redistribution of CX43 from gap junctions at intercalated disks to sites subjacent to the lateral sarcolemma (so-called lateralization) have been described in myocardium from DMD patients and *mdx* mice (reviewed in (17)). A recent study in *mdx* mice showed that CX43 is involved in the pathogenesis of DMD cardiomyopathy and that hypophosphorylation at specific serine residues (S325/S328/S330) is related to changes in CX43 expression, distribution, and function in DMD cardiomyopathy (26). Dysregulation of CX43 expression in *DMD*^Y/-^ pigs provides an additional model for studying these mechanisms and answering the question if CX43 expression can be normalized upon therapeutic restoration of dystrophin, e.g. by correcting the *DMD* reading frame via CRISPR/Cas9 (27).

Particularly in the myocardium of advanced aged *DMD*^Y/-^ pigs (and *DMD*^+/-^ carriers), a notable infiltration of eosinophils was observed, accompanying the typical DMD-associated multifocal polyphasic histopathological lesions. Extensive infiltration of eosinophils into areas of damaged myocardium has previously also been reported in human patients with myocardial infarction and in a corresponding pig model (28), as well as in the *mdx* mouse model, where eosinophils are involved in modulation of the inflammatory response, cardiac remodeling, and promotion of fibrosis (29, 30).

As observed in adolescent DMD patients (reviewed in (31)), 3-month-old *DMD* ^Y/-^ pigs showed impaired systolic heart function, i.e. a 19% (p < 0.001) reduced LV ejection fraction, which is consistent with early stages of dilatative cardiomyopathy (DCM). Additionally, LV fractional shortening, another diagnostic parameter for DMD-related DCM (32), was 27% (p < 0.01) smaller in *DMD* ^Y/-^ vs. WT pigs. Systolic cardiac dysfunction has also been described in the GRMD model, but only in adult animals between age 30 and 45 months (33). In contrast, the *DMD* ^Y/-^ pig model develops -in line with the accelerated disease phenotype - early stages of heart failure, a major cause of death in DMD, already at a young age, offering timely readouts of efficacy of new therapeutic concepts.

In addition to heart failure and progressive muscle weakness, many patients with DMD have cognitive and/or behavioral problems, which have great impact on the quality of life of both patients and their families (reviewed in (34)). To evaluate if *DMD* ^Y/-^ pigs resemble also these aspects of human DMD, we performed two behavioral tests which both revealed significant differences from WT. In particular, the Black and White Discrimination Test showed significantly impaired learning ability of *DMD* ^Y/-^ pigs. Molecular profiling of different brain regions available in our dedicated DMD biobank may provide mechanistic insights into the cognitive deficits associated with DMD.

A “byproduct” of generating *DMD* ^Y/-^ pigs by breeding are *DMD* ^+/-^ carrier pigs, which can be used for breeding but also for pathophysiological studies. In the present study, we concentrated on alterations of the heart since cardiac manifestations are present in a significant proportion of DMD carriers (35, 36).

Indeed, the 7-year-old *DMD* ^+/-^ founder sow #3040 showed severe pathological alterations of the heart, and 3/20 *DMD* ^+/-^ carriers showed macroscopic or microscopic alterations already at age 6 months. The biobank established from this collective is an interesting resource to screen for biomarkers that correlate with disease severity. We also showed that myocardial glucose utilization as detected by PET/CT is a promising parameter for identifying *DMD*^+/-^ carriers with cardiac abnormalities at an earlier age (3 months), which may be useful for longitudinal studies of disease progression in carriers or the translational validation of new therapeutic concepts. An example is targeted degradation of the nitric oxide synthase inhibitor asymmetric dimethylarginine (ADMA), which was recently shown to improve cardiac function and exercise tolerance in *mdx* carrier mice (37).

Many therapeutic approaches for DMD are in development aiming to restore dystrophin (e.g. gene replacement with microdystrophin, exon skipping, gene editing, stop-condon readthrough), utrophin upregulation, or targeting downstream effects dystrophin deficiency to preserve muscle tissue for as long as possible (reviewed in (1, 38)). Our scalable pig model for DMD allows comparative testing of different therapeutic and diagnostic strategies. Proof of princple has been provided by testing a CRISPR/Cas9-based approach to restore an intact reading frame in *DMD*Δ52 pigs (27) and by validating multispectral optoacoustic tomography (MSOT) as a new noninvasive imaging biomarker for assessing DMD disease progression (39). Importantly, groups of *DMD*^+/-^ carrier sows can be estrus-cycle synchronized and inseminated with semen from the same boar to reliably produce large cohorts of *DMD*^Y/-^ pigs and WT littermate controls for studies with statistically relevant group sizes, allowing for fast and effective testing of new treatment options in a large animal model for DMD with high clinical resemblance to the human phenotype and similar disease progression.

## Methods

### Generation of female DMD^+/-^ pigs and establishment of a breeding herd

Deletion of exon 52 of the porcine *DMD* gene was performed in principle as described in Klymiuk, et al. (10). In brief, the targeting bacterial artificial chromosome CH242-9G11 was modified to carry instead of exon 52 a neomycin resistance cassette. Subsequently, the modified BAC was nucleofected into a female primary kidney cell line (PKCf) of a German Landrace × Swabian-Hall hybrid background. Single-cell clones were generated according to Richter, et al. (40). From each cell clone genomic DNA was isolated, and cell clones with a heterozygous deletion of *DMD* exon 52 were identified by comparing its copy number with those of two reference loci (*POU5F1* and *NANOG*). Of 258 screened cell clones, 9 had one intact and one *DMD* allele lacking exon 52. One of these cell clones was used for somatic cell nuclear transfer (SCNT), and the reconstructed embryos were transferred laparoscopically to estrous cycle synchronized recipient gilts (41). A total of 14 embryo transfers resulted in 4 litters with 7 live heterozygous *DMD*Δ52 (*DMD*^+/-^) piglets. One of them (#3040) could be raised to adulthood and served as founder of a *DMD* pig breeding herd. Therefore, #3040 was inseminated with semen from wild-type German Landrace boars. For genotyping, PCR analyses of the offspring were performed using the primer pair 5’-TGC ACA ATG CTG GAG AAC CTC A-3’ / 5’-GTT CTG GCT TCT TGA TTG CTG G-3’ for the detection of the wild-type *DMD* allele and primer pair 5’-CAG CTG TGC TCG ACG TTG TC-3’ / 5’-GAA GAA CTC GTC AAG AAG GCG ATA G-3’ to detect the *DMD*Δ52 allele. In addition to the founder sow #3040, 6 F1 and 3 F2 *DMD*^+/-^ sows were inseminated with semen from WT boars to enlarge the breeding herd. Semen from a single *DMD*^Y/-^ boar (#6964) that could be raised to sexual maturity was used for insemination of 4 *DMD*^+/-^ sows (one of them founder #3040).

### Blood collection and clinical chemistry

Animals had free access to food and water before blood collection from the right jugular vein. Blood samples were taken using Serum Monovettes^®^ (Sarstedt, Nümbrecht). After clotting at room temperature for 30 min, serum was separated by centrifugation at 1.800 rcf for 20 min at 4 °C. The blood serum was stored at -80 °C until further processing (storage period did not exceed 6 months). CK values were determined using the CKL ACN 057 kit on a Cobas 311 Analyser System (Roche).

### Echocardiography

In order to evaluate the heart phenotype of the DMD pigs we performed a standard 2D transthoracic echocardiography (Esaote MyLab X8) in *DMD* ^Y/-^ (n = 4) and WT pigs (n = 7). All animals were born on the same day, raised in the same environment and examined at 3 months of age (102 or 103 days). The same person performed all echocardiographic examinations.

The pigs were kept under general anesthesia which was induced with 20 mg/kg ketamine (Ursotamin^®^, Serumwerke Bernburg) and 2 mg/kg azaperone (Azaporc^®^, Serumwerke Bernburg) and maintained with 4 mg/kg/h of propofol (Propofol 2%, Fresenius Kabi). Pigs were placed in right lateral recumbency.

Systolic function and dimensions of the heart were evaluated in the right parasternal long axis four chamber view using the Teichholz formula in M-mode and the Simpson method of discs in B-mode. The average of three measurements over different heart cycles was taken. To rule out pathologies influencing cardiac function and morphology a complete echocardiogram was performed.

### PET/CT analysis of myocardial glucose uptake

Positron emission tomography/computed tomography (PET/CT) data acquisition was performed in 3-month-old *DMD*^+/-^ carrier pigs (n = 5) and WT controls (n = 5). Two days before PET/CT examination, the animals were transported to the Walter Brendel Centre of Experimental Medicine (LMU Munich), where they were housed in individual cages in a temperature- and humidity-controlled room with feeding twice a day and free access to water.

For PET/CT acquisition, pigs were premedicated (intramuscular azaperone 2 mg/kg body weight, ketamine 20 mg/kg body weight) and transported to the Department of Nuclear Medicine (LMU Munich). In each ear, a catheter was placed in the *Vena auricularis caudalis*, one for radiotracer application and one for maintenance of anesthesia (xylazine 1 mg/kg body weight; ketamine 20 mg/kg body weight). The venous access was also used to measure the blood glucose (BG) level (Accu-Chek Performa blood glucose meter). Respiratory rate was monitored and logged every five minutes. Body temperature was measured (electronic clinical thermometer, reer GmbH) immediately before the PET/CT scan and 60 and 80 min after tracer injection. In addition, an electrocardiogram (ECG) lead according to Einthoven was carried out during the entire measurement period to monitor the animals and to enable an ECG-triggered (10 gates/cycle) PET data analysis (Ivy Model 7600, Ivy Biomedical Systems). The pigs were placed in supine position on a vacuum mattress (VacuSoft, UNGER Medizintechnik GmbH & Co.KG) to keep the animals in the same position on the PET/CT scanner for the whole time. The PET/CT procedure was performed on a Siemens Biograph 64 TruePoint PET/CT scanner (Siemens Healthcare GmbH). At the beginning, an x-ray topogram was run to determine the exact location of the heart. A CT scan was performed and used for attenuation correction of the PET data. Simultaneously with bolus injection of the glucose-analog 2-deoxy-2-[^18^F]fluoro-D-glucose ([^18^F]FDG; 293.65 ± 17.69 MBq), PET acquisition (one bed position of 17.6 cm axial extent) was started for 60 min. ECG-gated data in the time window 30 to 60 min p.i. were reconstructed (Siemens TrueX (3i21s)) including CT-based attenuation and model-based scatter correction (voxel size 2.67×2.67×3.0 mm^3^).

Image values were converted to standardized uptake values (SUV) by using the body weight and the injected activity. End-systolic myocardial data were volumetrically sampled, and polar maps of left ventricular myocardial SUV were generated by using a validated software (42, 43). After normalization to their maximum value, regional patterns of myocardial glucose uptake were evaluated using the 17-segment model of the American Society for Nuclear Cardiology (44). Mean values in the segments corresponding to the territories of the coronary arteries [right coronary artery (RCA), left circumflex artery (LCX), left anterior descending (LAD)] were determined for each dataset.

### Histopathology

All animals were euthanised over the ear vein. Tissue specimens of various skeletal muscles (including triceps brachii muscle, gluteobiceps muscle, diaphragm, and larynx muscle), of ventricular myocardium (each three transmural sections from the left, right and septal myocardium), and of the different parts of the gastrointestinal tract (GIT) were systematically sampled as described previously (45). Tissue samples were formalin fixed and routinely embedded in paraffin. Sections of 3 µm thickness were stained with hematoxylin-eosin. Giemsa-stained sections were prepared (46) for facilitated demonstration of degenerative and necrotic muscle fiber alterations and differentiation of inflammatory cells populations (macrophages, lymphocytes, plasma cells, eosinophils and neutrophils). Stained sections were examined with an Olympus BX 41 light microscope (Olympus). Digital images were acquired at 20-fold objective magnification using a digital camera (DP 72, Olympus) and adapted software (VIS-Visiopharm Integrator System™ Version 3.4.1.0, Visiopharm A/S). Masson’s Trichrome (MT) stain using a standard protocol (46) was performed of all GIT locations. Each section of the GIT was scored for intra- and submucosal accumulation of collagenous fibers as well as inflammatory mucosal alterations. Scoring system comprised the ranges 1 - 5 in accordance with the grade (absent, sub-significant, mild, moderate, severe) of fibrosis and inflammation, respectively. Pictures were taken using an orthoplan light microscope (851465, Leica Biosystems) and the pathoZoom software (Smart in Media GmbH).

### Immunohistochemistry

For immunohistochemical (IHC) detection of cellular DYS1 abundance patterns in paraffin sections of triceps muscle specimens, immunostaining was performed as described in Regensburger et al. 2019, using monoclonal mouse anti-DYS1 (Rod domain, no. NCL-DYS1; clone no. Dy4/6D3, Leica Biosystems, Wetzlar) as primary antibody, and biotinylated goat anti-mouse IgG (no. 115-065-146; Jackson ImmunoResearch, USA) as secondary antibody. In all IHC-experiments, appropriate negative control sections (omission of the first anti-DYS1-antibody) were used. Stained sections were photogaphed at 20-fold objective magnification using a digital camera (DP 72, Olympus) with adapted software (VIS-Visiopharm Integrator System™ Version 3.4.1.0, Visiopharm A/S)

### Immunofluorescence studies

Paraffin-embedded sections of 2-µm were prepared for histological analysis. In brief, sections were dewaxed, rehydrated in a descending alcohol series and subjected to antigen retrieval procedure using citrate buffer pH 6.0. Slides were washed with 1× PBS and blocked with 5% bovine serum albumin (BSA, Sigma-Aldrich) for 30 min at room temperature (RT) to saturate non-specific binding sites. Then, the samples were incubated with anti-connexin 43 (CX43, 1:100, Abcam, Ab3512) and and anti-cardiac troponin T (cTnT, 1:100, Abcam, Ab8295) antibodies diluted in 0.5% BSA solution overnight at 4 °C. After washing with 1× PBS, slides were incubated for 1 hour at 37 °C with fluorescent-conjugated secondary antibodies (1:1000, Thermo Fisher Scientific). Nuclei were counterstained with DAPI (1:1000) for 30 min and washed twice with 1× PBS. A Leica SP5 laser scanning confocal microscope (Leica Microsystem, Wetzlar, Germany) was used to acquire labelled samples. Analysis was performed in sequential scanning mode to rule out cross bleeding between channels, evaluating two cardiac areas of each animal (n = 3 per genotype and age group except for *DMD*^Y/-^ > 6 month: n = 2). Immunofluorescence images for CX43 were converted into black and white using ImageJ Software (47), according to the following sequence: Process>Binary>Convert to Binary. CX43 quantification, expressed as area of particles, was obtained using the command Analyze>Analyze Particles.

### Quantification of cardiac fibrosis

Paraffin-embedded sections (2-µm) were dewaxed and rehydrated in a descending alcohol series. After washing, the samples were stained with Masson’s Trichrome KIT reagents (Sigma-Aldrich, No.HT15, St. Louis, MO, USA) according to the manufacturer’s protocol. Four histological cardiac areas for each animal (n = 3 per genotype and age group except for *DMD*^Y/-^ > 6 month: n = 2) were evaluated to assess the size of the fibrotic area (blue staining) expressed as proportion (percentage) of the total area, using ImageJ Software.

### Sample preparation for proteome analysis

Frozen tissue samples from heart and skeletal muscle were transferred in a prechilled tubes and cryo-pulverized in a Covaris CP02 Automated Dry Pulverizer (Covaris Inc, MA, USA) according to the manufacturer’s instructions. Powdered tissue was lysed in 8 M urea/0.5 M NH4HCO3 by ultrasonicating (18 cycles of 10 s) using a Sonopuls HD3200 (Bandelin, Berlin, Germany). Pierce 660 nm Protein Assay (Thermo Fisher Scientific, Rockford, IL, USA) was used for protein quantification. 30 μg of protein was digested with Lys-C (FUJIFILM Wako Chemicals Europe GmbH, Germany) for 4 h and subsequently with modified porcine trypsin (Promega, WI, USA) for 16 h at 37 °C.

### Nano-LC-MS/MS analysis and bioinformatics

1.5 μg of peptide was injected on an UltiMate 3000 nano LC system online coupled to a Q-Exactive HF-X instrument (Thermo Scientific). Samples were transferred to a PepMap 100 C18 trap column (100 µm x 2 cm, 5 µM particles, Thermo Scientific) and separated on an analytical column (PepMap RSLC C18, 75 µm x 50 cm, 2 µm particles, Thermo Scientific) at 250 nL/min with a 160-min gradient of 3-25% of solvent B followed by 10-min increase to 40% and 5-min increase to 85%. Solvent A consisted of 0.1% FA in water and solvent B of 0.1% FA in acetonitrile. MS spectra were acquired using a top 15 data-dependent acquisition method on a Q Exactive HF-X mass spectrometer.

Protein identification was done using Maxquant (v.1.6.17.0) (48), using its built-in search engine Andromeda (49) and the NCBI RefSeq Sus scrofa database (v.7-5-2020). Reverse peptides, contaminants and identifications only by site were excluded from further analyses. Proteins having at least two peptides detected in at least three replicates of each condition were tested for differential expression using the MS-EmpiRe pipeline (50). To handle missing values, imputation was performed using the DEP package (51). Proteins having a Benjamini & Hochberg (BH) corrected p-value < 0.05 and a fold-change ≥ |1.3| were regarded as significant. Data visualization was performed using R (52) with ggplot2 (53), ggfortify (54) and ComplexHeatmap (55). Overrepresentation analysis for significantly altered proteins was performed using WebGestalt (56) and the functional categories “GO Biological Process nonRedundant” and “KEGG”. Benjamini & Hochberg (BH) method was used for multiple testing adjustment.

### Combined linkage disequilibrium and linkage analysis (cLDLA)

Genomic DNA was isolated from freshly docked tails of newborn *DMD*^Y/-^ piglets using the DNasy^®^ Blood & Tissue Kit (Qiagen). Genome-wide SNP genotyping was performed by TZF Poing-Grub with the PorcineSNP60 v2 Genotyping BeadChip (Illumina) featuring more than 64,000 SNPs that uniformly span the porcine genome. Markers of the SNP chip were remapped to the *Sus scrofa* reference genome assembly Sscrofa11.1 (57). During quality control, SNPs with an unknown position, a minor allele frequency lower than 0.025 and a call rate per marker lower than 0.9 were excluded. After quality control, 48,392 SNPs and 148 individuals with a genotype call rate higher than 0.95 remained for further analysis. Missing genotypes were imputed and haplotypes reconstructed using the software BEAGLE 5 (58). In total, 149 animals were genotyped including *DMD*^Y/-^ cases (n = 125) as well as their parents and ancestors (n = 24) in a complex pedigree structure. Genotyping of both parents allowed BEAGLE to perform high quality haplotyping of all *DMD*^Y/-^ cases and their parents in the 133 trio (sire-dam-case) cohortes and thus substantially improved subsequent mapping. A subset of n = 98 *DMD*^Y/-^ pigs was selected for genome-wide mapping to ensure that only animals which died spontaneously or had to be euthanized in a terminal condition were included. The genome-wide mapping of quantitative trait loci (QTL) was performed using a combined linkage disequilibrium and linkage analysis (cLDLA) approach, as reported in Meuwissen, et al. (59) and adapted by Medugorac, et al. (60) and Gehrke, et al. (61). First, a genomic relationship matrix (62) was estimated and used for prediction of random polygenic effects. Moreover, the genomic relationship matrix considered and corrected for complex family relationships in our mapping population. Second, for sliding windows consisting of 40 SNPs, identity by descent (IBD) probabilities for pairs of haplotypes (63) were estimated. The estimated haplotype IBD probabilities were then converted into a locus specific diplotype relationship matrix as suggested by Lee and Van der Werf (64). These locus specific diplotype relationship matrices were used for prediction of random QTL effects. The variance component analysis and likelihood estimation were performed in the mixed linear model with two random effects (QTL and polygenic) using the ASReml package (65). To estimate QTL effects at specific positions in the genome variance component analysis in the middle of each of the 40 SNPs windows sliding along genome was implemented. Using the logarithm of the likelihood estimated by ASReml for the model with (*log(L*_*1*_)) and without (*log(L*_*0*_)); corresponding to the null hypothesis) QTL locus effects, we calculated the likelihood-ratio test statistic (*LRT* = −2(*log(L*_*0*_) − *log(L*_*1*_)), which is known to be *χ*^*2*^ distributed with 1 degree of freedom. Appropriately, an *LRT* value higher than 11.0 was considered chromosome-wide significant (equivalent to Bonferroni corrected p < 0.05, i.e. p < 0.0009 divided by the average number of independent 40 SNP windows per chromosome). An LRT value higher than 16.5 was considered genome-wide significant. This threshold corresponds to a Bonferroni-corrected p < 0.05, i.e. p < 0.000049 divided by the number of genome-wide independent 40 SNP windows. Gehrke, et al. (61) demonstrated that *cLDLA* can be used for mapping of quantitative (as well as binary) traits. In the present study, the cLDLA mapping was performed with life expectancy as quantitative variable, i.e., length of life in days.

### Behavioral tests

We used 12 *DMD*^Y/-^ and 13 male WT littermates for the behavioral trials. The animals were from four litters and born at the same day. After weaning, all animals were kept in two genotypically mixed groups under standardized conditions. Training started at age 4 weeks. Trials started at age 5 weeks and ended at age 8 weeks. To standardize the experiments and to minimize all external influences, the training rounds for all animals took place on the same day and alternately for both genotypes. All animals were habituated under standardized conditions to human contact, short-term isolation and floor conditions in the start and trial boxes. The piglets were conditioned to receive banana juice from a blue dish. All experiments were performed in a trial box (3.0 × 1.8 m) in the same room, in which the animals were kept. During the trials, the animals were able to have acoustic but no visual contact. The floor of the trial box was completely perforated and cleaned between the rounds to avoid any influence of smells from previous rounds or animals.

Before starting the test, every animal was kept in the start box (0.5 × 1.8 m) for a minute to ensure uniform starting conditions. For both genotypes, trial numbers were assigned alternately. The animals always needed to pass the experiments in the same order. All experiments were recorded by a Canon EOS 550D camera.

A Novel Object Recognition Test was performed according to Lueptow (66) and Sondergaard, et al. (67). The animals were kept for 5 minutes in the experimental box with two objects with same structure but different shape and color. Red shades were avoided, because of the visible color spectrum of the pig’s eye. For this time frame the animals were able to explore both objects. Afterwards, they were returned to their group for 15 minutes. After this time, the animals were placed back in the experimental box with a new and an old object. The exploration time for each object was measured when the pigs nose was within a defined zone around the object (0.3 × 0.3 m). The exploration times of the new (T_new_) and of the old object (T_old_) as well as the total exploration time (T_total_ = T_new_ + T_old_) were recorded. The test was repeated after fifteen minutes, one day and one week.

A Black and White Discrimination Test was designed as a modified T maze test (68). Animals were placed in a trial box which was separated by a wall with two transparent doors (polystyrene; 0.25 × 0.4 m). The left door was marked with a black spot (Ø 10 cm) and the right door with a white spot (Ø 10 cm). Behind both doors, blue dishes with banana juice were visible. Before the trials, every piglet underwent two rounds of training to learn, that only through the black door they get access to the reward (banana juice). The first door the piglet chose was scored. Five rounds were performed this way. At the sixth round, the locations of both doors were switched, but still the black door offered access to the reward.

### Statistical analysis

Body weight data were analyzed using PROC MIXED (SAS 9.4; SAS Institute), taking the effects genotype, age, and interaction genotype×age as well as as well as the random effect of animal into account. Least squares means (LSMs) and standards errors (SEs) of LSMs were calculated for genotype×age and compared using student t-tests. Myocardial fibrosis and CX43 expression data as well as results of the Novel Object Recognition Test were statistically evaluated using PROC GLM (SAS 9.4), taking the effects of genotype, age/time, and interaction genotype×age/time into account. Results of the Black and White Discrimination Test were evaluates by chi-square analysis. Data from echocardiography were compared using student t-test. For fibrosis scores of the gastrointestinal tract, Mann-Whitney U test, one-sided was used. Differences between groups were considered as significant at p < 0.05; p-values are presented as: *: p < 0.05; **: p < 0.01; ***: p < 0.001.

## Study approval

All animal experiments were performed according to the German Animal Welfare Act and Directive 2010/63/EU on the protection of animals used for scientific purposes and were approved by the responsible animal welfare authority (Government of Upper Bavaria; permission 55.2-1-54-2532-163-2014).

## Author contributions

NK, BK, MK, HN, VZ, and EW developed the pig model; MS, LMF, AL, GW, SK, ABä, MHdA, CKu, and MCW performed the clinical and clinical-chemical characterization; ML, LMF, EK, SZ, and PB did the PET/CT studies; LMF, MS, CKa, AH, AL, BK, KM, and ABl established the biobanks; ABl, CM, LMF, MS, EK, KM, and RW performed pathological investigations; BS, FF, LAK, GJA, and TF did proteomics studies; MC, CB, and RR quantified myocardial fibrosis and CX43 expression; IM, LMF, and MS did the cLDLA mapping; MS, LMF, and EW wrote the manuscript; all authors edited and agreed to the final version of the manuscript.

## Acknowledgments

SNP genotypes of WT sires were provided by Martin Heudecker (EGZH, Poing-Grub) and Kay-Uwe Götz (ITZ, Poing-Grub). We would like to thank Harald Paul for excellent animal care; Christina Blechinger and Tatjana Schröter for excellent technical assistance; and Sophia Grotz, Melanie Janda, Stefanie Egerer, Isabel Hofmann and Georg Dhom for help with the tissue sampling. The project was supported by the Else Kröner-Fresenius Foundation (EKFS 2018_T20). In addition, this project received funding from the European Union’s Horizon 2020 research and innovation programme under the Marie Skłodowska-Curie grant agreement No 812660 (DohART-NET).

