## Supplemental figures and tables for "A scalable, clinically severe pig model for Duchenne muscular dystrophy"

*Resource and Technical Advance*

### Supplemental figures and tables

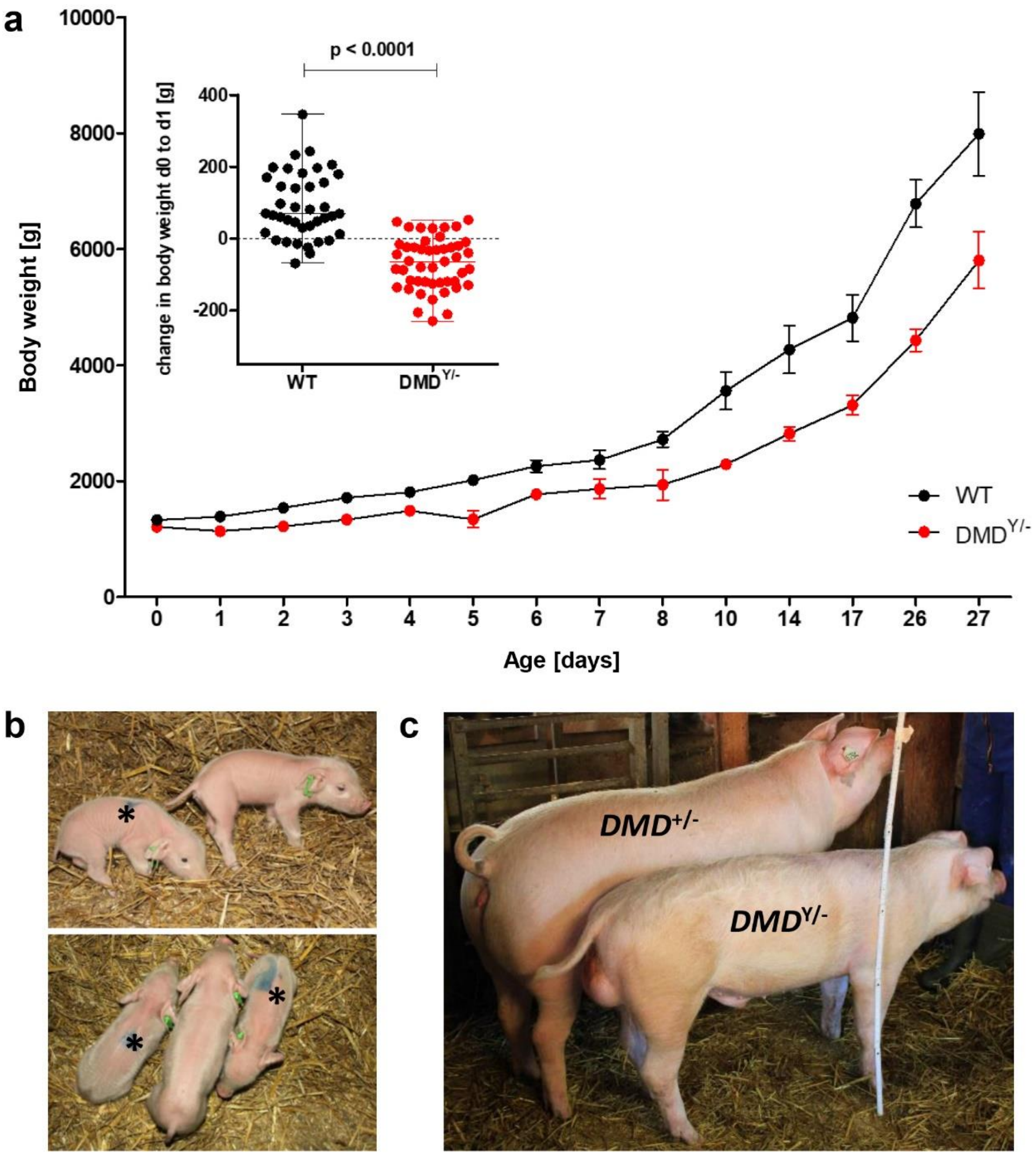


**Supplemental figure 1.** Reduced birth weight and weight gain of *DMD*$^{Y/-}$ piglets. **a**) Body weights of *DMD*$^{Y/-}$ pigs and male wild-type (WT) littermates in the first 27 days of life (n ≥ 16 per genotype until day 5; subsequently n ≥ 5 per genotype). For the period after day 8, data were adjusted by linear interpolation to the respective ages shown in the graph. Insert: Change in body weight of *DMD*$^{Y/-}$ piglets (n = 47) and male WT littermates (n = 39) between birth and age 24 hours. **b**) Representative pictures of 4-day-old *DMD*$^{Y/-}$ piglets (marked with asterisks) and a male WT littermate. **c**) Adolescent *DMD*$^{Y/-}$ pig #6790 and *DMD*$^{+/-}$ littermate at age 226 days.

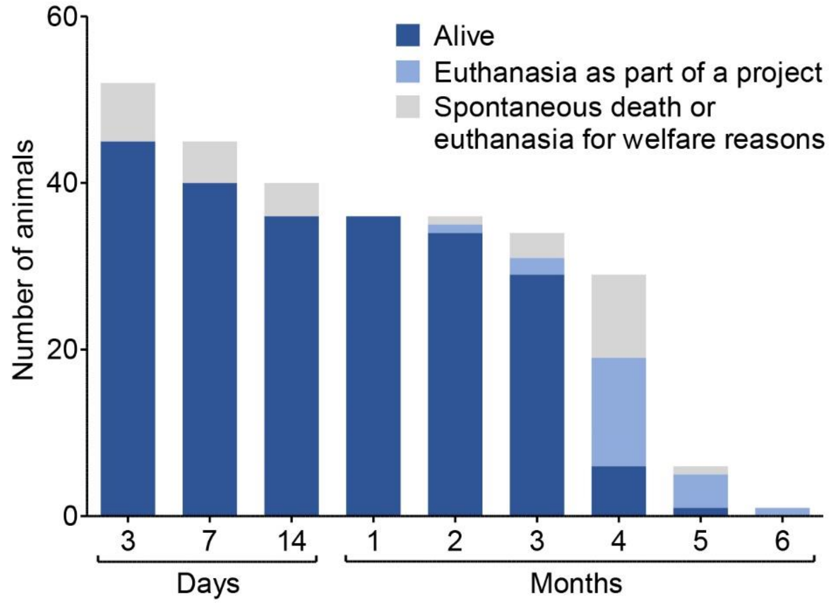


**Supplemental figure 2.** Reduced neonatal lethality of *DMD*$^{Y/-}$ piglets after optimisation of litter management (cohort of pigs born after 01/2020).

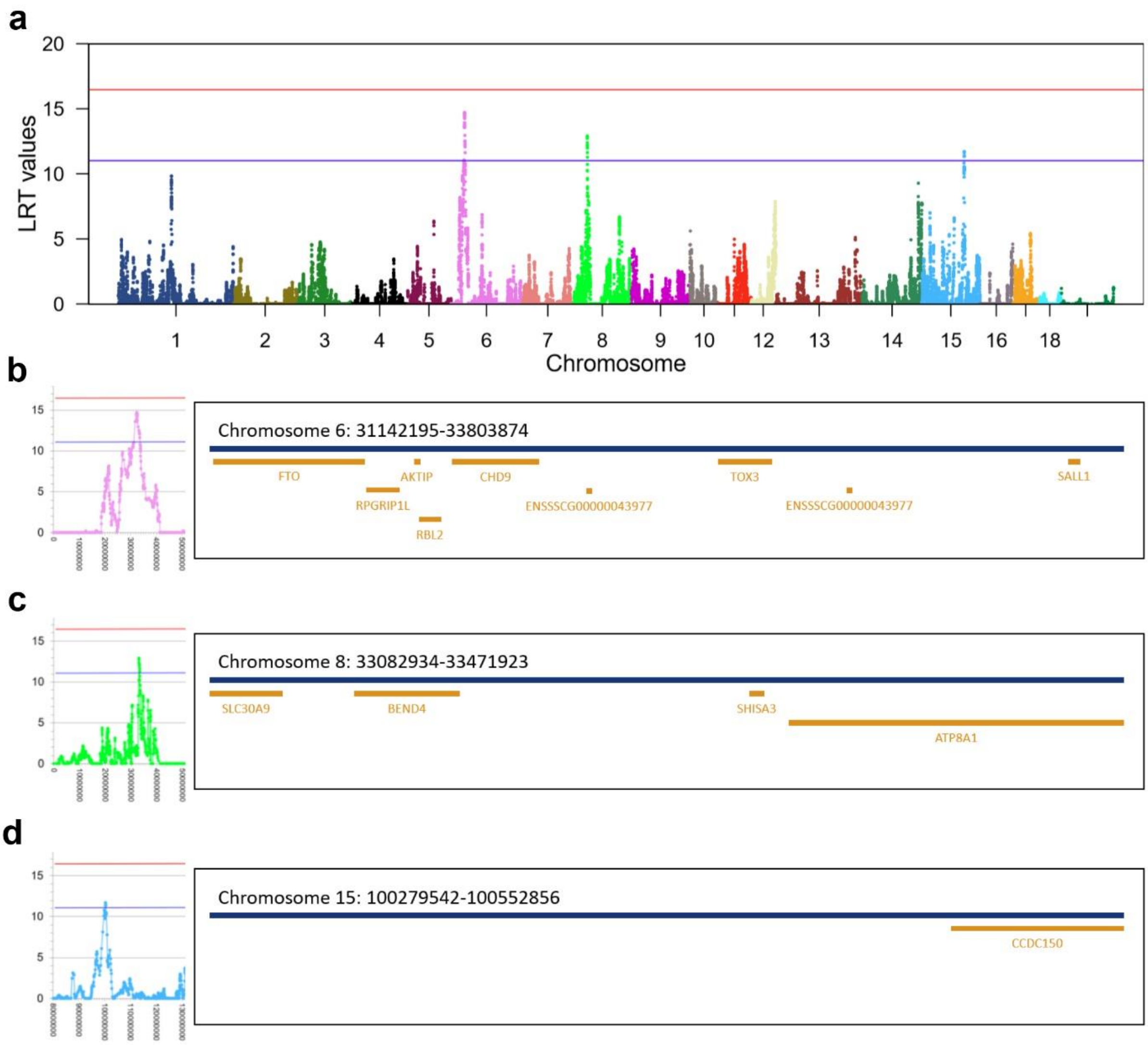


**Supplemental figure 3.** Mapping of quantitative trait loci affecting the life expectancy of *DMD*$^{Y/-}$ pigs. **a**) ) The association between haplotypes and phenotypes (life expectancy) is tested using the likelihood ratio test statistic (*LRT*). The *LRT* values are shown on the y-axis and the tested positions on the pig chromosomes on the x-axis. Blue horizontal line indicates the level of chromosome-wide significance, red line the level of genome-wide significance. **b-d**) QTL regions and protein-coding genes in these regions on chromosomes 6 (**b**), 8 (**c**), and 15 (**d**).

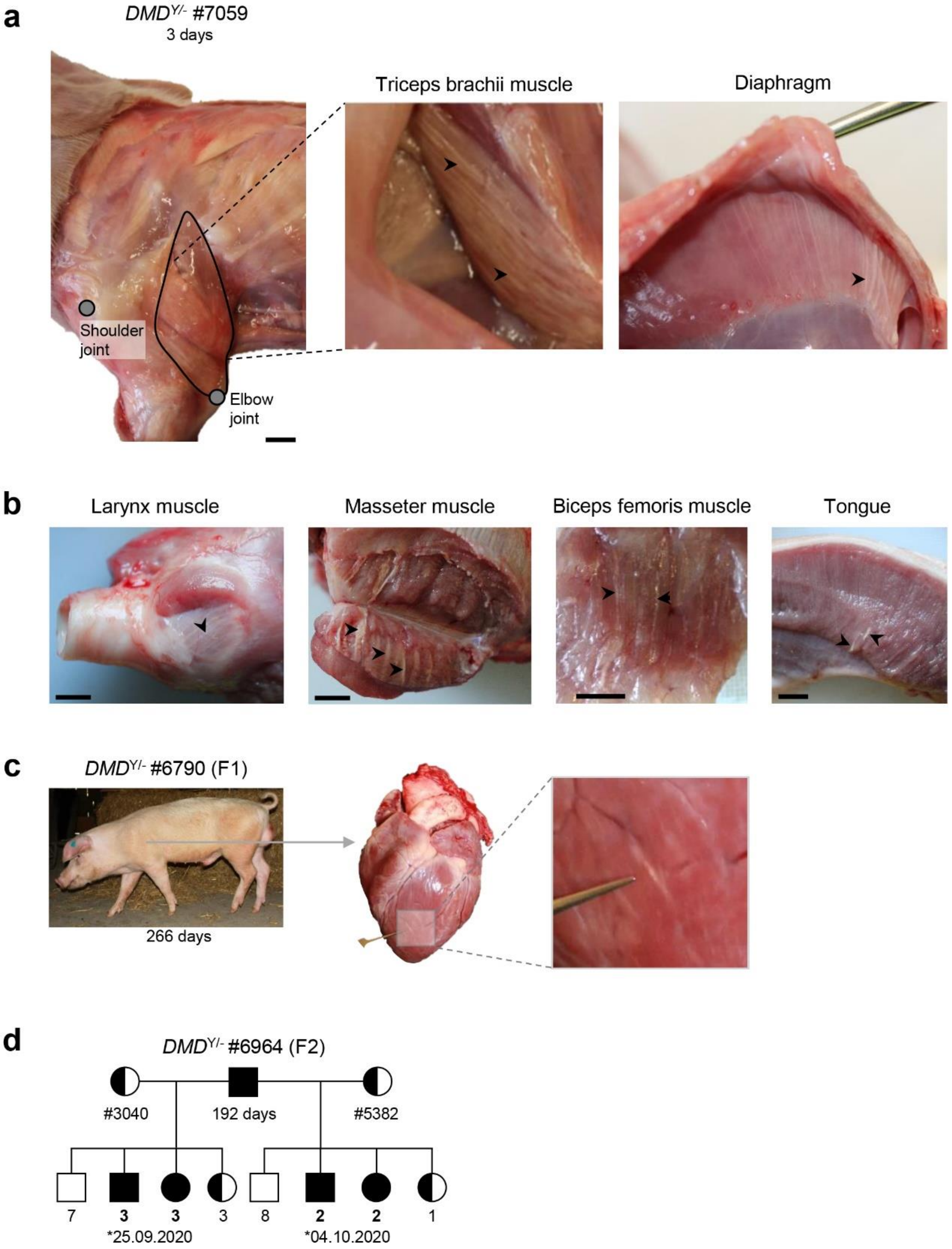


**Supplemental figure 4**. Macroscopic changes of skeletal muscle and heart in *DMD*$^{Y/-}$ pigs surviving less than one week (**a**), 3-4 months (**b**) or more than 6 months (**c**). **a**) Short-term survivors showed prominent macroscopic alterations of various skeletal muscles, such as streaky white muscle fiber degeneration. Scale bar = 1 cm. **b**) In *DMD*$^{Y/-}$ pigs surviving for 3-4 months, multiple skeletal muscles showed macroscopic signs of degeneration, whereas the heart was macroscopically inconspicuous. Scale bar = 1 cm. **c**) The longest surviving *DMD*$^{Y/-}$ boar #6790 displayed macroscopic myocardial lesions. **d**) From another long-term surviving *DMD*$^{Y/-}$ boar (#6964), semen could be recovered and used for insemination of *DMD*$^{+/-}$ carrier sows, which gave birth to offspring, including *DMD*$^{+/-}$ and *DMD*$^{-/-}$.

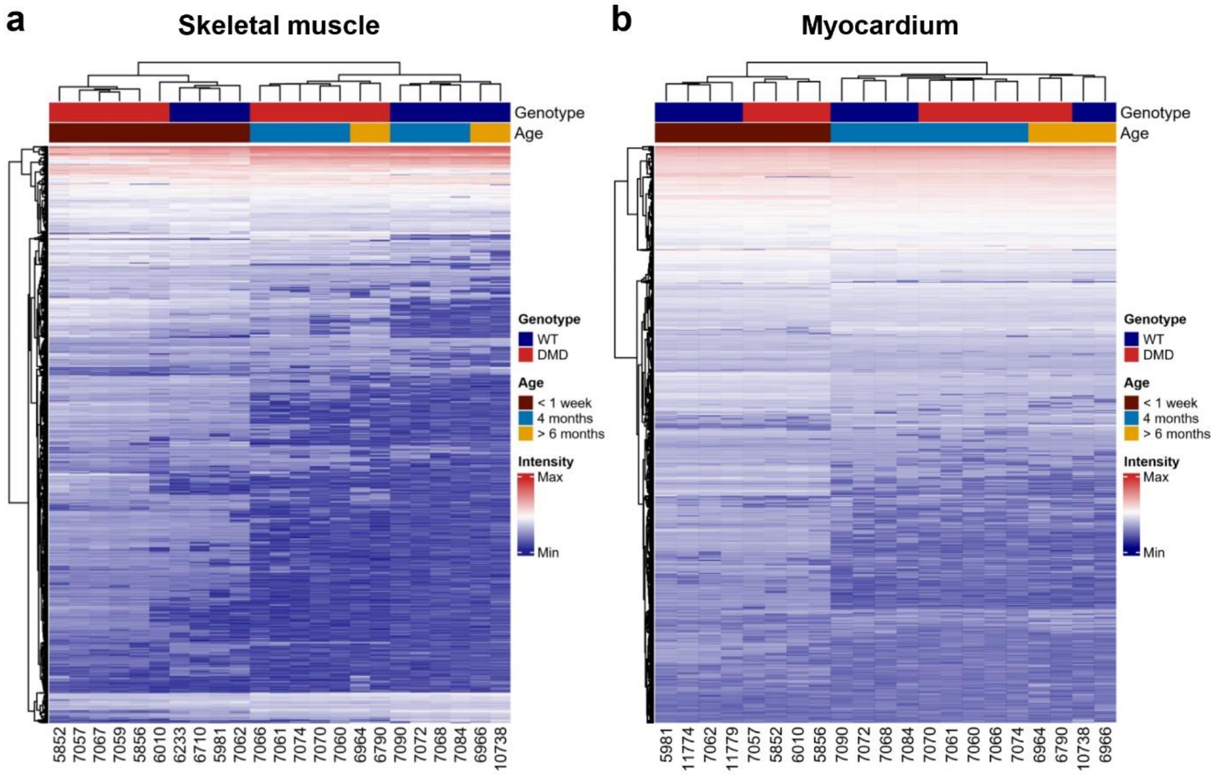


**Supplemental figure 5**. Unsupervised hierarchical clustering of normalized protein intensities from skeletal muscle (**a**) and myocardium (**b**) of < 1 week old, 4 months old, and > 6 months old WT and *DMD*$^{Y/-}$ animals**.**

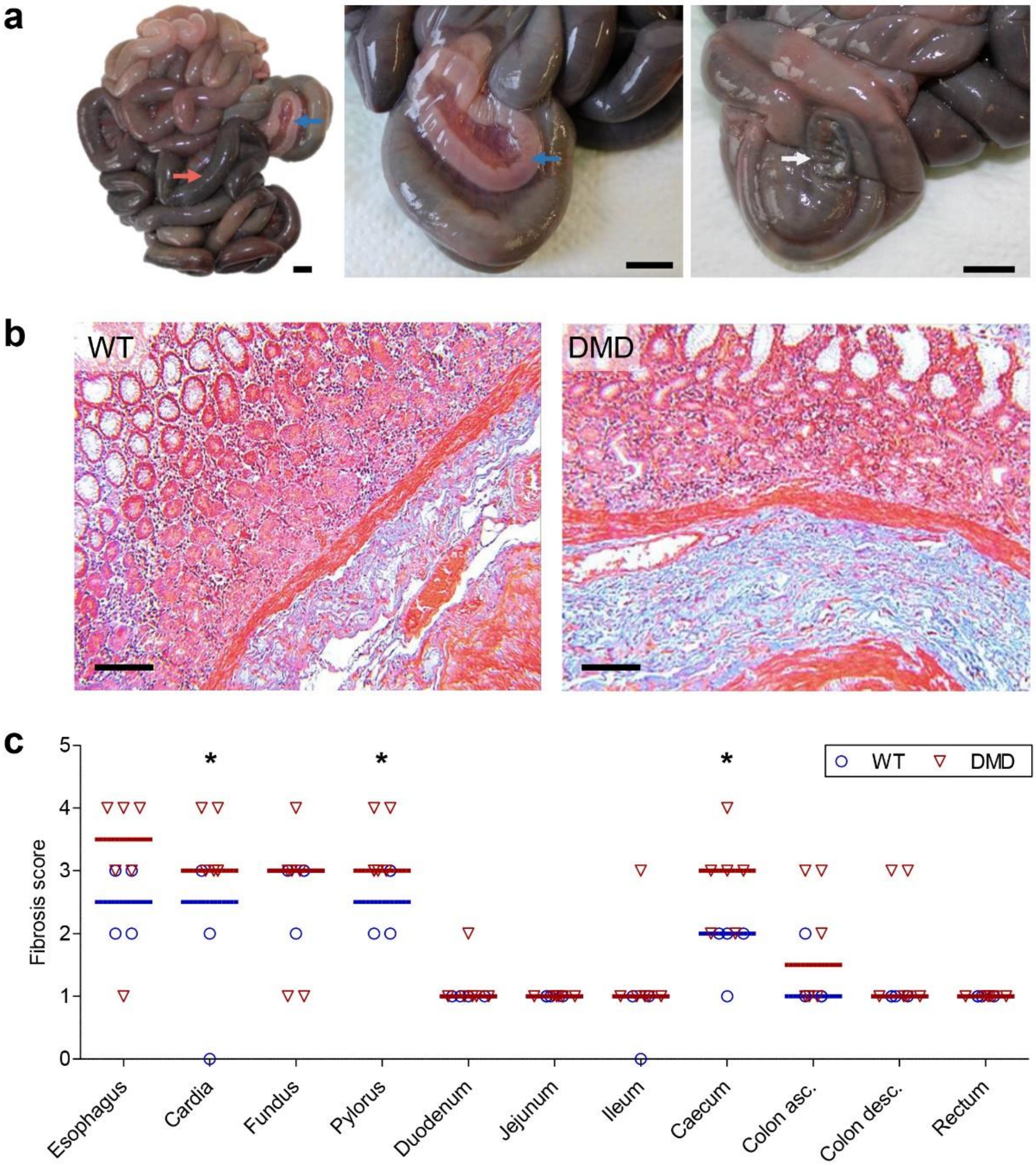


**Supplemental figure 6**. Macroscopic and histological alterations of the gastrointestinal tract in *DMD*$^{Y/-}$ pigs. **a**) Macroscopic findings in a 6-day-old *DMD*$^{Y/-}$ piglet showing clinical symptoms of a gastrointestinal phenotype. Obstruction of the large intestine (red arrow), empty intestine section subsequent to the obstructed part (blue arrow), peak of the ascending large intestine with porous site (white arrow). Bars = 1 cm. **b**) Histology of the stomach cardia from a 4-month-old wild-type (WT) and an age-matched *DMD*$^{Y/-}$ pig. The *DMD*$^{Y/-}$ pig shows an increased deposition of collagenous fibers (green-blue colored). Masson's Trichrome stain. Bar = 250 µm. **c**) Fibrosis scores in different parts of the gastrointestinal tract. 1 = absent; 2 = sub-significant; 3 = mild; 4 = moderate; 5 = severe. Asterisks indicate significant differences between *DMD*$^{Y/-}$ and WT pigs (Mann-Whitney U test, one-sided: $p < 0.05$).

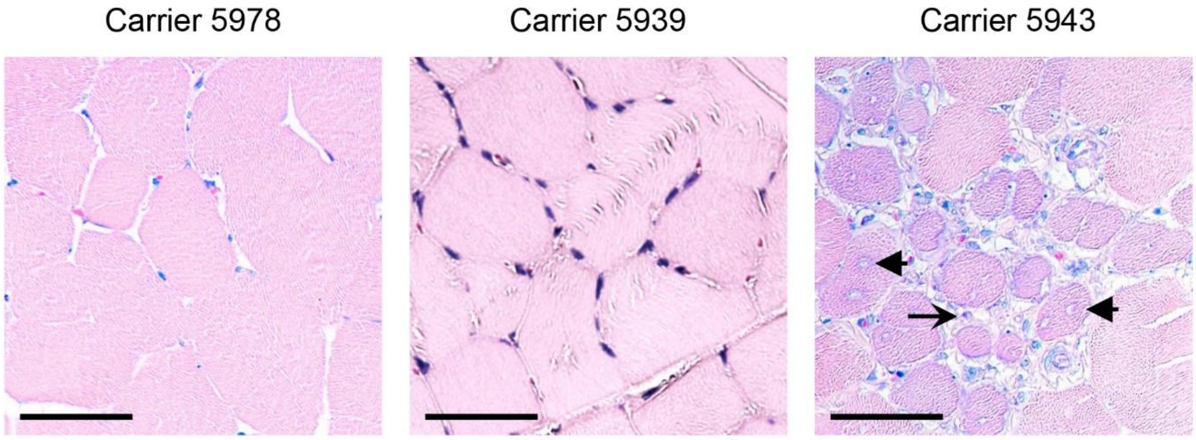


**Supplemental figure 7**. Variable histological alterations of skeletal muscle (triceps brachii) in 6-month-old *DMD*$^{+/-}$ carrier pigs. Histopathological alterations are indicated (arrows: inflammatory cell infiltration comprising of macrophages/histiocytes, plasma cells and few lymphocytes; arrowheads: degenerating skeletal myocytes with centralized nuclei). Paraffin sections, Giemsa-staining. Bars = 50 µm.

**Supplemental table 1.** Breeding of F1 and F2 *DMD*$^{+/-}$ offspring of founder sow #3040

| Gen. | Sow | M/F sow | Boar (WT) | Litter | Birth date | WT m | WT f | *DMD*$^{+/-}$ | *DMD*$^{Y/-}$ |
|---|---|---|---|---|---|---|---|---|---|
| F1 | **#5153** | #3040/Eposo | Isomer | 1 | 2017-10-18 | 1 | 6 | 4 | 5 |
| | | | Casanova | 2 | 2018-03-16 | 1 | 2 | 4 | 5 |
| | | | Boletto | 3 | 2018-08-09 | 7 | 4 | 2 | 3 |
| F1 | **#5381** | #3040/Rabatz | Costa | 1 | 2018-01-16 | 4 | 1 | 5 | 2 |
| | | | Costa | 2 | 2018-06-14 | 5 | 0 | 0 | 0 |
| | | | Costa | 3 | 2018-11-08 | 3 | 3 | 6 | 2 |
| F1 | **#5382** | #3040/Rabatz | Isomer | 1 | 2018-03-09 | 4 | 4 | 3 | 9 |
| | | | Boletto | 2 | 2018-08-03 | 1 | 6 | 7 | 4 |
| | | | Costa | 3 | 2019-03-27 | 4 | 8 | 1 | 3 |
| | | | Costa | 4 | 2019-08-22 | 5 | 1 | 5 | 5 |
| F1 | **#5383** | #3040/Rabatz | Costa | 1 | 2017-11-18 | 1 | 4 | 3 | 6 |
| | | | Costa | 2 | 2018-04-12 | 4 | 2 | 4 | 2 |
| | | | Casanova | 3 | 2018-09-06 | 3 | 5 | 5 | 3 |
| | | | Cor | 4 | 2019-10-08 | 3 | 2 | 7 | 2 |
| F1 | **#6314** | #3040/Costa | Rabart | 1 | 2019-12-02 | 4 | 3 | 3 | 3 |
| | | | Rabart | 2 | 2020-04-21 | 0 | 6 | 4 | 2 |
| | | | Last | 3 | 2021-02-19 | 4 | 5 | 3 | 2 |
| F1 | **#6794** | #3040/Cor | Rabart | 1 | 2020-08-24 | 5 | 1 | 3 | 5 |
| | | | Last | 2 | 2021-02-19 | 2 | 5 | 5 | 1 |
| F2 | **#6225** | #5382/Boletto | Cor | 1 | 2019-09-02 | 0 | 1 | 2 | 1 |
| | | | Last | 2 | 2020-07-02 | 4 | 2 | 5 | 5 |
| | | | Unox | 3 | 2020-11-27 | 3 | 3 | 5 | 3 |
| F2 | **#6243** | #5153/Boletto | Cor | 1 | 2019-06-13 | 5 | 3 | 3 | 2 |
| | | | Rabart | 2 | 2020-04-16 | 3 | 0 | 5 | 7 |
| | | | Last | 3 | 2020-09-18 | 5 | 4 | 3 | 2 |
| | | | Last | 4 | 2021-02-19 | 5 | 3 | 4 | 3 |
| F2 | **#6245** | #5153/Boletto | Rabart | 1 | 2019-06-19 | 3 | 4 | 3 | 3 |
| | | | Rabart | 2 | 2019-11-09 | 2 | 3 | 4 | 7 |
| | | | Rabal | 3 | 2020-04-02 | 1 | 0 | 3 | 6 |
| | | | Rabart | 4 | 2020-09-05 | 4 | 2 | 3 | 3 |
| | | | Last | 5 | 2021-02-19 | 2 | 5 | 4 | 8 |
| **Total** | | | | **31** | | **98** | **98** | **118** | **114** |

M/F sow = mother/father of the sow; WT m = wild-type male; WT f = wild-type female.

| Supplemental table 2. Overview of samples collected in the DMD-Biobank | | | |
|---|---|---|---|
| Organ system | Organ/tissue | Location/tissue compartment | Samples |
| Central nervous system | Brain | Hippocampus | FFPE, -80°C |
| | | Temporo-ventral body of hippocampus | FFPE, -80°C |
| | | Amygdala | FFPE, -80°C |
| | | Cerebellar cortex | FFPE, -80°C |
| Cardiovascular system | Heart transmural (epicardium, myocardium, endocardium) | Left ventricle basis | Cryo, FFPE, -80°C |
| | | Left ventricle middle | Cryo, FFPE, -80°C |
| | | Left ventricle apex | Cryo, FFPE, -80°C |
| | | Septum basis | Cryo, FFPE, -80°C |
| | | Septum middle | Cryo, FFPE, -80°C |
| | | Septum apex | Cryo, FFPE, -80°C |
| | | Right ventricle basis | Cryo, FFPE, -80°C |
| | | Right ventricle middle | Cryo, FFPE, -80°C |
| | | Right ventricle apex | Cryo, FFPE, -80°C |
| | | Right atrial appendage | FFPE, -80°C |
| | | Left atrial appendage | FFPE, -80°C |
| Musculo-skeletal system | Triceps brachii | Central muscle head | Cryo, FFPE, -80°C |
| | Gluteobiceps | Central muscle head | Cryo, FFPE, -80°C |
| | Diaphragm | From the left thorax wall | Cryo, FFPE, -80°C |
| Gastro-intestinal tract | Tongue | Basis/middle/apex | FFPE |
| | Esophagus | | Cryo, FFPE, -80°C |
| | Stomach | Cardiac region | Cryo, FFPE, -80°C |
| | | Fundus region | Cryo, FFPE, -80°C |
| | | Pylorus region | Cryo, FFPE, -80°C |
| | Duodenum | | Cryo, FFPE, -80°C |
| | Jejunum | | Cryo, FFPE, -80°C |
| | Ileum | | Cryo, FFPE, -80°C |
| | Caecum | | Cryo, FFPE, -80°C |
| | Ascending colon | | Cryo, FFPE, -80°C |
| | Descending colon | | Cryo, FFPE, -80°C |
| | Rectum | | Cryo, FFPE, -80°C |
| Hepato-pancreatic system | Liver | | Cryo, FFPE, -80°C |
| Uro-genital system | Testis | Left testis | Cryo, FFPE |
| | Epididymis | Head/Body/Tail | FFPE |
| | Kidney | Cortex | Cryo, FFPE, -80°C |
| | | Medulla | Cryo, FFPE, -80°C |
| Respiratory tract | Larynx | | FFPE |
| | Lung | | Cryo, FFPE, -80°C |
| Hematopoietic system | Spleen | | Cryo, FFPE, -80°C |
| Body fluids | Blood serum | | -80°C |
| | Li-heparin blood plasma | | -80°C |
| | EDTA blood plasma | | -80°C |

| Supplemental table 4. Overview of samples collected in the Carrier-Biobank | | | |
|---|---|---|---|
| **Organ system** | **Organ/tissue** | **Location/tissue compartment** | **Samples** |
| **Cardiovascular system** | **Heart transmural (epicardium, myocardium, endocardium)** | **Left ventricle** | **Cryo, FFPE, -80°C** |
| | | **Septum** | **Cryo, FFPE, -80°C** |
| | | **Right ventricle** | **Cryo, FFPE, -80°C** |
| **Musculo-skeletal system** | **Triceps brachii** | **Central muscle head** | **Cryo, FFPE, -80°C** |
| | **Gluteobiceps** | **Central muscle head** | **Cryo, FFPE, -80°C** |
| | **Diaphragm** | **From the left thorax wall** | **Cryo, FFPE, -80°C** |
| | **Longissimus dorsi** | **Central muscle head** | **Cryo, FFPE, -80°C** |
| **Body fluids** | **Blood serum** | | **-80°C** |
| | **Li-heparin blood plasma** | | **-80°C** |
| | **EDTA blood plasma** | | **-80°C** |
| | **Urine** | | **-80°C** |

**Supplemental table 5.** Haplotypes/diplotypes at the quantitative trait locus on chromosome 6 affecting life expectancy of DMD pigs.

PhC = Phenotype (life expectancy in days); DipC = Diplotype; DipE = estimated effect of diplotype

Haplotypes/diplotypes with positive and negative effects are shaded in green and red, respectively.

| Nr | TierLID | mHap\|_____._____\|_____X_____\|_____._____\|_____0 | pHap\|_____._____\|_____X_____\|_____._____\|_____0 | HapC (m) | HapC (p) | PhC | DipC | DipE | PolygenE |
|---|---|---|---|---|---|---|---|---|---|
| 44 | DMD6828 | AGAAGAACCGGAAAGAAGAGGCGGGGGAAGAAAGGAGGAC | GAACACGAAGGAAAGGGAAGAAGGGGAAAAGGAGGGAGGC | 3 | 17 | 9 | 26 | -40 | 0,000010 |
| 32 | DMD6293 | AGAAGAACCGGAAAGAAGAGGCGGGGGAAGAAAGGAGGAC | GACAGCGACGGAGAGGAGGAGAGGGGAAAGAAAGGGAAGC | 3 | 14 | 2 | 20 | -36 | -0,000010 |
| 30 | DMD6235 | AGAAAAGACGGAGAAGGAGAGAGGGGAAAGAAGAGGGAAC | GAAAGAAAAGGAGGGAAAGGGAAAAAAGGAGGGAGAAAAA | 13 | 5 | 6 | 19 | -33 | -0,000030 |
| 31 | DMD6238 | GAAAGAAAAGGAGGGAAAGGGAAAAAAGGAGGGAGAAAAA | AGAAAAGACGGAGAAGGAGAGAGGGGAAAGAAGAGGGAAC | 5 | 13 | 3 | 19 | -33 | -0,000030 |
| 53 | DMD7057 | GAAAGAAAAGGAGGGAAAGGACAGGAGAAAGGAGGGAAGC | GAAAGAAAAGGAGGGAAAGGGAAAAAAGGAGGGAGAAAAA | 8 | 5 | 5 | 30 | -15 | -0,000030 |
| 55 | DMD7059 | GAAAGAAAAGGAGGGAAAGGGAAAAAAGGAGGGAGAAAAA | AGAAGAAAAGGAGGGAAAGGACAGGAGAAAGGAGGGAAGC | 5 | 19 | 3 | 32 | -15 | -0,000020 |
| 46 | DMD6831 | GACAGAACAGAAGGGGAAGGACAAAAGAAGAAAGGGAGGC | GAACACGAAGGAAAGGGAAGAAGGGGAAAAGGAGGGAGGC | 6 | 17 | 25 | 27 | -7,6 | -0,000010 |
| 47 | DMD6836 | GAACACGAAGGAAAGGGAAGAAGGGGAAAAGGAGGGAGGC | GACAGAACAGAAGGGGAAGGACAAAAGAAGAAAGGGAGGC | 17 | 6 | 6 | 27 | -7,6 | -0,000020 |
| 48 | DMD6837 | GACAGAACAGAAGGGGAAGGACAAAAGAAGAAAGGGAGGC | GAACACGAAGGAAAGGGAAGAAGGGGAAAAGGAGGGAGGC | 6 | 17 | 4 | 27 | -7,6 | 0,000000 |
| 42 | DMD6755 | GACAGAACAGAAGGGGAAGGACAAAAGAAGAAAGGGAGGC | GAAAGAAAAGGAGGGAAAGGGAAAAAAGGAGGGAGAAAAA | 6 | 5 | 24 | 24 | -4,6 | -0,000010 |
| 75 | DMD7258 | GACAGAACAGAAGGGGAAGGACAAAAGAAGAAAGGGAGGC | GAAAGAAAAGGAGGGAAAGGGAAAAAAGGAGGGAGAAAAA | 6 | 5 | 94 | 24 | -4,6 | 0,000010 |
| 10 | DMD5721 | GACAGAACAGAAGGGGAAGGACAAAAGAAGAAAGGGAGGC | AGAAGAACCGGAAAGAAGAGGCGGGGGAAGAAAGGAGGAC | 6 | 3 | 4 | 6 | -3,3 | 0,000010 |
| 15 | DMD5852 | AGAAGAACCGGAAAGAAGAGGCGGGGGAAGAAAGGAGGAC | GACAGAACAGAAGGGGAAGGACAAAAGAAGAAAGGGAGGC | 3 | 6 | 3 | 6 | -3,3 | 0,000000 |
| 16 | DMD5856 | GACAGAACAGAAGGGGAAGGACAAAAGAAGAAAGGGAGGC | AGAAGAACCGGAAAGAAGAGGCGGGGGAAGAAAGGAGGAC | 6 | 3 | 3 | 6 | -3,3 | 0,000000 |
| 22 | DMD6011 | GACAGAACAGAAGGGGAAGGACAAAAGAAGAAAGGGAGGC | AGAAGAACCGGAAAGAAGAGGCGGGGGAAGAAAGGAGGAC | 6 | 3 | 50 | 6 | -3,3 | 0,000030 |
| 37 | DMD6577 | AGAAGAACCGGAAAGAAGAGGCGGGGGAAGAAAGGAGGAC | AGAAAAAAAAGGAGGAAAGGACAAAAGAAGAAAGGGAGGC | 3 | 15 | 71 | 22 | -3,1 | 0,000000 |
| 4 | DMD4059 | GACAGAACAGAAGGGGAAGGACAAAAGAAGAAGAGGAAAC | AGAAGAACCGGAAAGAAGAGGCGGGGGAAGAAAGGAGGAC | 4 | 3 | 114 | 3 | -2,8 | 0,000010 |
| 27 | DMD6214 | AGAAAAGACGGAGAAGGAGAGAGGGGAAAGAAGAGGGAAC | GACAGAACAGAAGGGGAAGGACAAAAGAAGAAAGGGAGGC | 13 | 6 | 38 | 17 | 0,9 | -0,000020 |
| 28 | DMD6215 | GACAGAACAGAAGGGGAAGGACAAAAGAAGAAAGGGAGGC | AGAAAAGACGGAGAAGGAGAGAGGGGAAAGAAGAGGGAAC | 6 | 13 | 3 | 17 | 0,9 | -0,000040 |
| 11 | DMD5724 | AGAAAAAAAAGGAGGAAAGAGCAAGGAAAGAAAGGGAAAC | AGAAGAACCGGAAAGAAGAGGCGGGGGAAGAAAGGAGGAC | 7 | 3 | 139 | 7 | 2,55 | 0,000050 |
| 21 | DMD5984 | AGAAAAAAAAGGAGGAAAGAGCAAGGAAAGAAAGGGAAAC | AGAAGAACCGGAAAGAAGAGGCGGGGGAAGAAAGGAGGAC | 7 | 3 | 102 | 7 | 2,55 | 0,000040 |
| 24 | DMD6016 | AGAAGAACCGGAAAGAAGAGGCGGGGGAAGAAAGGAGGAC | AGAAAAAAAAGGAGGAAAGAGCAAGGAAAGAAAGGGAAAC | 3 | 7 | 54 | 7 | 2,55 | 0,000020 |
| 33 | DMD6311 | AGAAGAACCGGAAAGAAGAGGCGGGGGAAGAAAGGAGGAC | AGAAAAAAAAGGAGGAAAGAGCAAGGAAAGAAAGGGAAAC | 3 | 7 | 3 | 7 | 2,55 | 0,000000 |
| 35 | DMD6411 | AGAAAAAAAAGGAGGAAAGAGCAAGGAAAGAAAGGGAAAC | AGAAGAACCGGAAAGAAGAGGCGGGGGAAGAAAGGAGGAC | 7 | 3 | 121 | 7 | 2,55 | 0,000020 |
| 45 | DMD6829 | AGAAAAAAAAGGAGGAAAGAGCAAGGAAAGAAAGGGAAAC | AGAAGAACCGGAAAGAAGAGGCGGGGGAAGAAAGGAGGAC | 7 | 3 | 1 | 7 | 2,55 | -0,000030 |
| 8 | DMD5139 | GAAAGAAAAGGAGGGAAAGGGAAAAAAGGAGGGAGAAAAA | GAACAAGCCAGAAGGGAAGAGAAGGGAAAAGGAGAAAAAA | 5 | 2 | 69 | 5 | 3,89 | 0,000010 |
| 9 | DMD5145 | GAACAAGCCAGAAGGGAAGAGAAGGGAAAAGGAGAAAAAA | GAAAGAAAAGGAGGGAAAGGGAAAAAAGGAGGGAGAAAAA | 2 | 5 | 5 | 5 | 3,89 | -0,000020 |
| 19 | DMD5970 | GACAGCGACGGAGAGGAGGAGAGGGGAAAGAAAGGAGGAC | GAACAAGCCAGAAGGGAAGAGAAGGGAAAAGGAGAAAAAA | 12 | 2 | 2 | 13 | 4,87 | -0,000020 |
| 80 | DMD7310 | AGAAGAACCGGAAAGAAGAGGCGGGGGAAGAAAGGAGGAC | GAACAAGCCAGAAGGGAAGAGAAGGGAAAAGGAGAAAAAA | 3 | 2 | 5 | 37 | 5,27 | -0,000020 |
| 3 | DMD4058 | GAACACGAAGGGGAAGGAGGACAGGGGAAGAAAGGAAAAC | AGAAGAACCGGAAAGAAGAGGCGGGGGAAGAAAGGAGGAC | 1 | 3 | 24 | 2 | 7,81 | -0,000020 |
| 5 | DMD4070 | AGAAGAACCGGAAAGAAGAGGCGGGGGAAGAAAGGAGGAC | GAACACGAAGGGGAAGGAGGACAGGGGAAGAAAGGAAAAC | 3 | 1 | 23 | 2 | 7,81 | -0,000020 |
| 29 | DMD6231 | GAAAGAAAAGGAGGGAAAGGGAAAAAAGGAGGGAGAAAAA | AGAAAAGACGGAGAAGGAGGACAGGGGAAGAAAGGAAAAC | 5 | 11 | 71 | 18 | 8,3 | -0,000050 |
| 39 | DMD6707 | GAAAGAAAAGGAGGGAAAGGGAAAAAAGGAGGGAGAAAAA | AGAAAAGACGGAGAAGGAGGACAGGGGAAGAAAGGAAAAC | 5 | 11 | 5 | 18 | 8,3 | -0,000030 |
| 40 | DMD6711 | GAAAGAAAAGGAGGGAAAGGGAAAAAAGGAGGGAGAAAAA | AGAAAAGACGGAGAAGGAGGACAGGGGAAGAAAGGAAAAC | 5 | 11 | 26 | 18 | 8,3 | -0,000030 |
| 60 | DMD7069 | AGAAAAGACGGAGAAGGAGGACAGGGGAAGAAAGGAAAAC | GAAAGAAAAGGAGGGAAAGGGAAAAAAGGAGGGAGAAAAA | 11 | 5 | 10 | 18 | 8,3 | -0,000060 |
| 62 | DMD7074 | GAAAGAAAAGGAGGGAAAGGGAAAAAAGGAGGGAGAAAAA | AGAAAAGACGGAGAAGGAGGACAGGGGAAGAAAGGAAAAC | 5 | 11 | 116 | 18 | 8,3 | -0,000020 |
| 77 | DMD7282 | GAAAGAAAAGGAGGGAAAGGGAAAAAAGGAGGGAGAAAAA | AGAAAAGACGGAGAAGGAGGACAGGGGAAGAAAGGAAAAC | 5 | 11 | 39 | 18 | 8,3 | 0,000000 |
| 89 | DMD7467 | AGAAAAGACGGAGAAGGAGGACAGGGGAAGAAAGGAAAAC | GAAAGAAAAGGAGGGAAAGGGAAAAAAGGAGGGAGAAAAA | 11 | 5 | 68 | 18 | 8,3 | 0,000020 |
| 90 | DMD7469 | GAAAGAAAAGGAGGGAAAGGGAAAAAAGGAGGGAGAAAAA | AGAAAAGACGGAGAAGGAGGACAGGGGAAGAAAGGAAAAC | 5 | 11 | 3 | 18 | 8,3 | 0,000020 |
| 91 | DMD7470 | GAAAGAAAAGGAGGGAAAGGGAAAAAAGGAGGGAGAAAAA | AGAAAAGACGGAGAAGGAGGACAGGGGAAGAAAGGAAAAC | 5 | 11 | 95 | 18 | 8,3 | 0,000030 |
| 93 | DMD7472 | GAAAGAAAAGGAGGGAAAGGGAAAAAAGGAGGGAGAAAAA | AGAAAAGACGGAGAAGGAGGACAGGGGAAGAAAGGAAAAC | 5 | 11 | 115 | 18 | 8,3 | 0,000040 |
| 95 | DMD7486 | GAAAGAAAAGGAGGGAAAGGGAAAAAAGGAGGGAGAAAAA | AGAAAAGACGGAGAAGGAGGACAGGGGAAGAAAGGAAAAC | 5 | 11 | 86 | 18 | 8,3 | 0,000000 |
| 18 | DMD5937 | AGAAGAACCGGAAAGAAGAGGCGGGGGAAGAAAGGAGGAC | AGAAAAGACGGAGAAGGAGGACAGGGGAAGAAAGGAAAAC | 3 | 11 | 77 | 12 | 9,63 | 0,000020 |
| 25 | DMD6212 | AGAAGAACCGGAAAGAAGAGGCGGGGGAAGAAAGGAGGAC | AGAAAAGACGGAGAAGGAGGACAGGGGAAGAAAGGAAAAC | 3 | 11 | 5 | 12 | 9,63 | -0,000060 |
| 78 | DMD7306 | AGAAAAGACGGAGAAGGAGGACAGGGGAAGAAAGGAAAAC | AGAAGAACCGGAAAGAAGAGGCGGGGGAAGAAAGGAGGAC | 11 | 3 | 2 | 12 | 9,63 | -0,000020 |
| 81 | DMD7320 | AGAAAAGACGGAGAAGGAGGACAGGGGAAGAAAGGAAAAC | AGAAGAACCGGAAAGAAGAGGCGGGGGAAGAAAGGAGGAC | 11 | 3 | 56 | 12 | 9,63 | -0,000010 |
| 17 | DMD5936 | GACAGAACAGAAGGGGAGAGGCGGGGGAAGAAAGGAGGAC | AGAAAAGACGGAGAAGGAGGACAGGGGAAGAAAGGAAAAC | 10 | 11 | 136 | 11 | 12,3 | 0,000030 |
| 49 | DMD6859 | GAACACGAAGGGGAGAAAGGACAGGGGAAGAAAGGAAAAC | AGAAAAGACGGAGAAGGAGAGAGGGGAAAGAAGAGGGAAC | 18 | 13 | 84 | 28 | 12,4 | 0,000020 |
| 65 | DMD7169 | AGAAAAGACGGAGAAGGAGAGAGGGGAAAGAAGAGGGAAC | AGAAAAGACGGAGAAGGAGGACAGGGGAAGAAAGGAAAAC | 13 | 11 | 109 | 35 | 13,9 | 0,000030 |
| 68 | DMD7173 | AGAAAAGACGGAGAAGGAGGACAGGGGAAGAAAGGAAAAC | AGAAAAGACGGAGAAGGAGAGAGGGGAAAGAAGAGGGAAC | 11 | 13 | 118 | 35 | 13,9 | 0,000040 |
| 85 | DMD7390 | AGAAAAGACGGAGAAGGAGAGAGGGGAAAGAAGAGGGAAC | AGAAAAGACGGAGAAGGAGGACAGGGGAAGAAAGGAAAAC | 13 | 11 | 5 | 35 | 13,9 | -0,000020 |
| 12 | DMD5725 | GACAGAACAGAAGGGGAAGGACAAAAGAAGAAAGGGAGGC | GAAAGAAAAGGAGGGAAAGGACAGGAGAAAGGAGGGAAGC | 6 | 8 | 107 | 8 | 18 | 0,000030 |
| 20 | DMD5983 | GAAAGAAAAGGAGGGAAAGGACAGGAGAAAGGAGGGAAGC | AGAAAAAAAAGGAGGAAAGAGCAAGGAAAGAAAGGGAAAC | 8 | 7 | 123 | 14 | 23,7 | 0,000040 |
| 51 | DMD6967 | AGAAAAAAAAGGAGGAAAGAGCAAGGAAAGAAAGGGAAAC | GAAAGAAAAGGAGGGAAAGGACAGGAGAAAGGAGGGAAGC | 7 | 8 | 46 | 14 | 23,7 | -0,000020 |
| 34 | DMD6409 | GACAGAACAGAAGGGGAAGGACAAAAGAAGAAAGGGAGGC | GACAGAACAGAAGGGGAAGGACAAAAGAAGAAAGGGAGGC | 6 | 6 | 75 | 21 | 28,8 | 0,000010 |
| 56 | DMD7060 | AGAAAAGACGGAGAAGGAGGACAGGGGAAGAAAGGAAAAC | GAAAGAAAAGGAGGGAAAGGACAGGAGAAAGGAGGGAAGC | 11 | 8 | 124 | 33 | 30,9 | 0,000000 |
| 71 | DMD7230 | AGAAAAGACGGAGAAGGAGGACAGGGGAAGAAAGGAAAAC | GAAAGAAAAGGAGGGAAAGGACAGGAGAAAGGAGGGAAGC | 11 | 8 | 95 | 33 | 30,9 | 0,000030 |
| 73 | DMD7232 | GAAAGAAAAGGAGGGAAAGGACAGGAGAAAGGAGGGAAGC | AGAAAAGACGGAGAAGGAGGACAGGGGAAGAAAGGAAAAC | 8 | 11 | 2 | 33 | 30,9 | -0,000010 |
| 38 | DMD6649 | GACAGAACAGAAGGGGAAGGACAAAAGAAGAAAGGAGGAC | AGAAAAAAAAGGAGGAAAGAGCAAGGAAAGAAAGGGAAAC | 16 | 7 | 96 | 23 | 34,7 | 0,000020 |
| 13 | DMD5763 | AGAAGAACCGGAAAGAAAGAGAAGGGAAAAGGAGAAAAAA | GACAGAACAGAAGGGGAAGGACAAAAGAAGAAAGGGAGGC | 9 | 6 | 82 | 9 | 36,3 | 0,000000 |
| 14 | DMD5769 | GAACAAGCCAGAAGGGAAGAGAAGGGAAAAGGAGAAAAAA | GACAGAACAGAAGGGGAAGGACAAAAGAAGAAAGGGAGGC | 2 | 6 | 108 | 10 | 37,4 | 0,000010 |
| 36 | DMD6575 | GACAGAACAGAAGGGGAAGGACAAAAGAAGAAAGGGAGGC | GAACAAGCCAGAAGGGAAGAGAAGGGAAAAGGAGAAAAAA | 6 | 2 | 123 | 10 | 37,4 | 0,000020 |
| 70 | DMD7228 | GAACAAGCCAGAAGGGAAGAGAAGGGAAAAGGAGAAAAAA | GACAGAACAGAAGGGGAAGGACAAAAGAAGAAAGGGAGGC | 2 | 6 | 125 | 10 | 37,4 | 0,000030 |
| 72 | DMD7231 | GAACAAGCCAGAAGGGAAGAGAAGGGAAAAGGAGAAAAAA | GACAGAACAGAAGGGGAAGGACAAAAGAAGAAAGGGAGGC | 2 | 6 | 74 | 10 | 37,4 | 0,000010 |
| 74 | DMD7233 | GACAGAACAGAAGGGGAAGGACAAAAGAAGAAAGGGAGGC | GAACAAGCCAGAAGGGAAGAGAAGGGAAAAGGAGAAAAAA | 6 | 2 | 111 | 10 | 37,4 | 0,000030 |
| 82 | DMD7327 | GAACAAGCCAGAAGGGAAGAGAAGGGAAAAGGAGAAAAAA | GACAGAACAGAAGGGGAAGGACAAAAGAAGAAAGGGAGGC | 2 | 6 | 116 | 10 | 37,4 | 0,000010 |
| 7 | DMD4796 | GACAGAACAGAAGGGGAAGGACAAAAGAAGAAGAGGAAAC | GAACAAGCCAGAAGGGAAGAGAAGGGAAAAGGAGAAAAAA | 4 | 2 | 97 | 4 | 37,7 | -0,000010 |
| 26 | DMD6213 | GACAGAACAGAAGGGGAAGGACAAAAGAAGAAAGGGAGGC | AGAAAAGACGGAGAAGGAGGACAGGGGAAGAAAGGAAAAC | 6 | 11 | 3 | 16 | 41,7 | -0,000050 |
| 41 | DMD6744 | AGAAAAGACGGAGAAGGAGGACAGGGGAAGAAAGGAAAAC | GACAGAACAGAAGGGGAAGGACAAAAGAAGAAAGGGAGGC | 11 | 6 | 6 | 16 | 41,7 | -0,000020 |
| 66 | DMD7170 | AGAAAAGACGGAGAAGGAGGACAGGGGAAGAAAGGAAAAC | GACAGAACAGAAGGGGAAGGACAAAAGAAGAAAGGGAGGC | 11 | 6 | 98 | 16 | 41,7 | 0,000020 |
| 67 | DMD7171 | GACAGAACAGAAGGGGAAGGACAAAAGAAGAAAGGGAGGC | AGAAAAGACGGAGAAGGAGGACAGGGGAAGAAAGGAAAAC | 6 | 11 | 98 | 16 | 41,7 | 0,000020 |
| 69 | DMD7176 | AGAAAAGACGGAGAAGGAGGACAGGGGAAGAAAGGAAAAC | GACAGAACAGAAGGGGAAGGACAAAAGAAGAAAGGGAGGC | 11 | 6 | 105 | 16 | 41,7 | 0,000020 |
| 76 | DMD7259 | AGAAAAGACGGAGAAGGAGGACAGGGGAAGAAAGGAAAAC | GACAGAACAGAAGGGGAAGGACAAAAGAAGAAAGGGAGGC | 11 | 6 | 192 | 16 | 41,7 | 0,000030 |
| 83 | DMD7387 | AGAAAAGACGGAGAAGGAGGACAGGGGAAGAAAGGAAAAC | GACAGAACAGAAGGGGAAGGACAAAAGAAGAAAGGGAGGC | 11 | 6 | 118 | 16 | 41,7 | 0,000000 |
| 84 | DMD7389 | GACAGAACAGAAGGGGAAGGACAAAAGAAGAAAGGGAGGC | AGAAAAGACGGAGAAGGAGGACAGGGGAAGAAAGGAAAAC | 6 | 11 | 118 | 16 | 41,7 | 0,000000 |
| 23 | DMD6012 | GAACAAGCCAGAAGGGAAGAGAAGGGAAAAGGAGAAAAAA | AGAAAAAAAAGGAGGAAAGAGCAAGGAAAGAAAGGGAAAC | 2 | 7 | 80 | 15 | 43,1 | 0,000000 |
| 79 | DMD7307 | GAACAAGCCAGAAGGGAAGAGAAGGGAAAAGGAGAAAAAA | GAACAAGCCAGAAGGGAAGAGAAGGGAAAAGGAGAAAAAA | 2 | 2 | 11 | 36 | 45,8 | -0,000040 |
| 64 | DMD7082 | AGAAAAAAAAGGAGGAAAGAGCAAGGAAAGAAAGAAAAAA | AGAAAAGACGGAGAAGGAGGACAGGGGAAGAAAGGAAAAC | 20 | 11 | 118 | 34 | 47,3 | -0,000010 |
| 52 | DMD6975 | AGAAAAAAAAGGAGGAAAGAGCAAGGAAAGAAAGGGAAAC | AGAAAAGACGGAGAAGGAGGACAGGGGAAGAAAGGAAAAC | 7 | 11 | 5 | 29 | 47,4 | -0,000030 |
| 97 | DMD7500 | AGAAAAGACGGAGAAGGAGGACAGGGGAAGAAAGGAAAAC | AGAAAAAAAAGGAGGAAAGAGCAAGGAAAGAAAGGGAAAC | 11 | 7 | 137 | 29 | 47,4 | 0,000050 |
| 98 | DMD7503 | AGAAAAAAAAGGAGGAAAGAGCAAGGAAAGAAAGGGAAAC | AGAAAAGACGGAGAAGGAGGACAGGGGAAGAAAGGAAAAC | 7 | 11 | 72 | 29 | 47,4 | 0,000020 |
| 1 | DMD4055 | GAACACGAAGGGGAAGGAGGACAGGGGAAGAAAGGAAAAC | GAACAAGCCAGAAGGGAAGAGAAGGGAAAAGGAGAAAAAA | 1 | 2 | 66 | 1 | 48,4 | -0,000020 |
| 2 | DMD4056 | GAACACGAAGGGGAAGGAGGACAGGGGAAGAAAGGAAAAC | GAACAAGCCAGAAGGGAAGAGAAGGGAAAAGGAGAAAAAA | 1 | 2 | 104 | 1 | 48,4 | -0,000010 |
| 6 | DMD4794 | GAACAAGCCAGAAGGGAAGAGAAGGGAAAAGGAGAAAAAA | GAACACGAAGGGGAAGGAGGACAGGGGAAGAAAGGAAAAC | 2 | 1 | 97 | 1 | 48,4 | -0,000010 |
| 43 | DMD6790 | AGAAAAGACGGAGAAGGAGGACAGGGGAAGAAAGGAAAAC | GAACAAGCCAGAAGGGAAGAGAAGGGAAAAGGAGAAAAAA | 11 | 2 | 266 | 25 | 50,4 | 0,000060 |
| 50 | DMD6964 | GAACAAGCCAGAAGGGAAGAGAAGGGAAAAGGAGAAAAAA | AGAAAAGACGGAGAAGGAGGACAGGGGAAGAAAGGAAAAC | 2 | 11 | 192 | 25 | 50,4 | 0,000000 |
| 63 | DMD7081 | AGAAAAGACGGAGAAGGAGGACAGGGGAAGAAAGGAAAAC | GAACAAGCCAGAAGGGAAGAGAAGGGAAAAGGAGAAAAAA | 11 | 2 | 114 | 25 | 50,4 | -0,000010 |
| 86 | DMD7439 | GAACAAGCCAGAAGGGAAGAGAAGGGAAAAGGAGAAAAAA | AGAAAAGACGGAGAAGGAGGACAGGGGAAGAAAGGAAAAC | 2 | 11 | 112 | 25 | 50,4 | 0,000040 |
| 54 | DMD7058 | AGAAAAGACGGAGAAGGAGGACAGGGGAAGAAAGGAAAAC | AGAAAAGACGGAGAAGGAGGACAGGGGAAGAAAGGAAAAC | 11 | 11 | 109 | 31 | 54,6 | -0,000010 |
| 57 | DMD7061 | AGAAAAGACGGAGAAGGAGGACAGGGGAAGAAAGGAAAAC | AGAAAAGACGGAGAAGGAGGACAGGGGAAGAAAGGAAAAC | 11 | 11 | 124 | 31 | 54,6 | -0,000010 |
| 58 | DMD7066 | AGAAAAGACGGAGAAGGAGGACAGGGGAAGAAAGGAAAAC | AGAAAAGACGGAGAAGGAGGACAGGGGAAGAAAGGAAAAC | 11 | 11 | 116 | 31 | 54,6 | -0,000030 |
| 59 | DMD7067 | AGAAAAGACGGAGAAGGAGGACAGGGGAAGAAAGGAAAAC | AGAAAAGACGGAGAAGGAGGACAGGGGAAGAAAGGAAAAC | 11 | 11 | 4 | 31 | 54,6 | -0,000070 |
| 61 | DMD7070 | AGAAAAGACGGAGAAGGAGGACAGGGGAAGAAAGGAAAAC | AGAAAAGACGGAGAAGGAGGACAGGGGAAGAAAGGAAAAC | 11 | 11 | 116 | 31 | 54,6 | -0,000030 |
| 87 | DMD7465 | AGAAAAGACGGAGAAGGAGGACAGGGGAAGAAAGGAAAAC | AGAAAAGACGGAGAAGGAGGACAGGGGAAGAAAGGAAAAC | 11 | 11 | 90 | 31 | 54,6 | 0,000020 |
| 88 | DMD7466 | AGAAAAGACGGAGAAGGAGGACAGGGGAAGAAAGGAAAAC | AGAAAAGACGGAGAAGGAGGACAGGGGAAGAAAGGAAAAC | 11 | 11 | 112 | 31 | 54,6 | 0,000020 |
| 92 | DMD7471 | AGAAAAGACGGAGAAGGAGGACAGGGGAAGAAAGGAAAAC | AGAAAAGACGGAGAAGGAGGACAGGGGAAGAAAGGAAAAC | 11 | 11 | 116 | 31 | 54,6 | 0,000030 |
| 94 | DMD7474 | AGAAAAGACGGAGAAGGAGGACAGGGGAAGAAAGGAAAAC | AGAAAAGACGGAGAAGGAGGACAGGGGAAGAAAGGAAAAC | 11 | 11 | 134 | 31 | 54,6 | 0,000040 |
| 96 | DMD7488 | AGAAAAGACGGAGAAGGAGGACAGGGGAAGAAAGGAAAAC | AGAAAAGACGGAGAAGGAGGACAGGGGAAGAAAGGAAAAC | 11 | 11 | 81 | 31 | 54,6 | -0,000010 |
